# Dietary fructose promotes MASH/HCC progression through enhanced intestinal HIF-2α-dependent iron absorption

**DOI:** 10.64898/2026.06.07.729655

**Authors:** Robert A. Mitchell, Manman Xu, Elizabeth Hudson, Madison S. Teer, Bradford G. Hill, Craig J. McClain, Ming Song

## Abstract

Dietary fructose is a major risk factor driving the progression of metabolic dysfunction-associated steatohepatitis (MASH) and hepatocellular carcinoma (HCC). However, the underlying fructose-induced nutrient-sensing pathway remains unclear. Here, we report that fructose facilitates iron absorption through the (Ketohexokinase) KHK/PKM2/HIF-2α axis, driving MASH and HCC development. Fructose aberrantly stabilizes intestinal HIF-α; this effect is abrogated by a KHK inhibitor and genetic *Khk* deletion. Mechanistically, fructose-induced metabolic reprogramming drives glutamine-dependent oxidative phosphorylation, leading to HIF-α stabilization, which is mediated by pyruvate kinase M2 (PKM2). A selective PKM2 inhibitor rescues reduced intestinal HIF-α stability in *Khk*-deficient mice. Furthermore, dietary fructose increases plasma iron levels. Conversely, *Khk*-deficient mice exhibit spontaneous systemic iron deficiency, characterized by hypochromic anemia. Moreover, *Khk* deficiency inhibits iron absorption in a HIF-2α-dependent manner. Finally, fructose promotes MASH and HCC progression in an iron-dependent manner. This study reveals a unique, therapeutically targetable nutrient-sensing pathway utilized by dietary fructose.

**In brief:** Mitchell et al. demonstrate that fructose consumption increases plasma iron levels, while KHK deficiency inhibits iron absorption in a HIF-2α-dependent manner. Mechanistically, dietary fructose-induced metabolic reprogramming aberrantly stabilizes intestinal HIF-α, which is mediated by PKM2. Fructose promotes MASH and HCC progression in an iron-dependent manner.

**Highlights:**

- Fructose aberrantly stabilizes intestinal HIF-α
- KHK is required for intestinal HIF-α stability
- KHK deficiency inhibits iron absorption in a HIF-2α-dependent manner
- Dietary fructose promotes MASH and HCC progression in an iron-dependent manner

## INTRODUCTION

Increased consumption of sugar-sweetened beverages (SSBs) has emerged as an important risk factor for obesity, type 2 diabetes mellitus (T2DM), metabolic dysfunction-associated steatotic liver disease (MASLD) and cancer. ^1,2^ The primary sweetener in SSBs is high-fructose corn syrup (HFCS), which is composed of almost equal amounts of fructose and glucose. ^3^ It is well known that fructose and glucose play divergent roles in the development of metabolic disorders, with fructose inducing more pronounced metabolic phenotypes, such as liver toxicity. ^4,5^

The liver was previously considered to be the primary site of fructose metabolism until recent studies showed that, in mice, fructose is primarily metabolized in the small intestine, as demonstrated by a stable isotope approach. Furthermore, fructose absorption and metabolism in the small intestine is saturable. Excess fructose is metabolized by the liver and gut microbiota. ^6,7^ Fructose metabolism in the small intestine (SI) reduces the load of ingested fructose reaching the liver; therefore, it was proposed that fructose metabolism in the SI protects against liver toxicity, ^6,7^ whereas the metabolic effects of fructose in the liver promote hepatic steatosis. ^8^ However, recent studies suggest the metabolic effects of fructose in the intestine are context-dependent, with one study showing that its metabolism in enterocytes facilitates lipid absorption and obesity when consumed with a high-fat diet. ^9^ Fructose metabolism by gut microbiota generates acetate, which serves as a substrate for *de novo* lipogenesis (DNL) via ACSS2-mediated acetyl-CoA. ^10^ On the other hand, fructose metabolism by enterocytes leads to the downregulation of tight-junction proteins (TJPs) and gut barrier dysfunction through endoplasmic-reticulum (ER) stress, inducing endotoxemia and hepatic inflammation, which promotes DNL. ^11^ Collectively, the integrated effects of fructose metabolism by the host and microbiota contribute to the pathogenesis of MASLD via the gut-liver axis.

Several lines of evidence demonstrate that fructose metabolism under hypoxia conditions is beneficial to cell survival and vital organ function ^9,12,13^. Unlike glucose, fructose is metabolized by KHK, leading to the generation of fructose 1-phosphate (F1P), which bypasses feedback inhibition of glycolysis via phosphofructokinase, thereby supporting survival under hypoxia. ^13^ Supporting this, fructose—but not glucose—metabolism enhances enterocytes survival and protects against apoptosis under hypoxia, leading to prolonged survival of enterocytes and villus hypertrophy. This, consequently, results in increased lipid absorption and exacerbated MASLD when consumed with a high-fat diet. ^9^ Mechanistically, HIF-1α drives alternative splicing that switches KHK-A to KHK-C through splice factor 3b subunit 1 (SF3B1), dictating fructose metabolism when oxygen supply is insufficient, ^12^ thus underpinning a molecular link between fructose metabolism and hypoxia.

Overall, fructose is preferentially used as a fuel under hypoxia conditions. ^9,12,13^ Physiologic hypoxia is a key feature of maintaining a healthy gut and hypoxia inducible factors (HIF-1α; HIF-2α) serve as important transcriptional regulators involved in the digestive system’s adaptation to low oxygen tensions (hypoxia). ^14,15^ Interestingly, HFCS feeding induces high expression of intestinal GLUT5 and KHK in mice, which is associated with upregulation of HIF-1α target protein, enolase-1 (ENO1) and lactate dehydrogenase A (LDHA), ^9^ suggesting a potential link between KHK and HIF signaling.

Intestinal HIF-2α signaling is essential for the maintenance of systemic iron homeostasis through transcriptional regulation of genes involved in iron absorption, including Dmt1. ^16,17^ Moreover, hepatic hepcidin crosstalk with intestine HIF-2α controls iron absorption under both physiologic and iron overload conditions. ^18^ Activation of intestinal HIF-2α signaling promotes hepatic steatosis, while intestinal-specific inhibition of HIF-2α signaling ameliorates it. ^19^ However, it is unknown whether HIF-2α-dependent iron absorption contributes to NASH and HCC development, and no links have ever been reported between dietary fructose consumption and intestinal HIF-2α stability. Fructose is primarily metabolized by KHK, leading to the generation of F1P in intestinal epithelial cells, ^6,9^ and F1P has recently been shown to specifically inhibit pyruvate kinase M2 (PKM2)-dependent catalytic activity. ^9^ Moreover, a PKM2 activator reduces intestinal iron absorption and prevents hepatic iron overload in a β-thalassemia mouse model. ^20^ Although PKM2 has a non-catalytic function that reportedly serves as a coactivator for HIF-dependent transactivation ^21^, the catalytic function for PKM2 in coordinating fructose/KHK-dependent effects on HIF-α stabilization and iron absorption remains elusive.

Here, we show that dietary fructose and glucose differentially regulate intestine HIF-α stability using HIF-α reporter (ODD-*luc*) mice. ^22^ While glucose markedly decreases intestinal HIF-α stability, fructose preserves it. Moreover, the “physiologic hypoxia” of the intestine exhibits apparent regional differences, being highest in the jejunum and lowest in the colon, which is positively associated with KHK expression in the intestine. By crossing ODD-*luc* mice ^22^ with *Khk*^+/+^ and *Khk*^-/-^ mice, we generated *Khk*^+/+^/ODD-*luc* and *Khk*^-/-^/ODD-*luc* mice, respectively, and discovered that KHK is required for intestinal HIF-α stability. Reduced HIF-α stability in *Khk*^-/-^ mice is rescued by a selective PKM2 inhibitor. We further show that fructose feeding increases plasma iron levels, and KHK deficiency inhibits iron absorption in a HIF-2α dependent manner. Consistent with this, *Khk*^-/-^ mice spontaneously develop hypochromic anemia, suggesting systemic iron deficiency. ^23^ Lastly, we show fructose promotes MASH and HCC progression in an iron-dependent manner. Collectively, our study identifies a unique nutrient-sensing pathway activated by dietary fructose that alters iron homeostasis through the KHK/HIF-2α axis.

## RESULTS

### Fructose aberrantly stabilizes intestinal HIF-α

Dietary fructose and glucose play differential roles in driving the development of MASLD. ^4,5^ However, the underlying mechanisms are poorly understood. Because glucose reportedly affects HIF-α stability ^24^, we first asked whether intestinal HIF-α stability was generally regulated by dietary fructose. To test this, we fed ODD-*luc* mice ^22^ with 15% (w/v) fructose (F), glucose (G) or a 1:1 mixture of fructose/glucose (FG) via drinking water *ad libitum* for 2 weeks. *Ex vivo* bioluminescence imaging (BLI) showed that mice fed fructose and the FG mixture exhibited significantly increased HIF-α stabilization (i.e., luminescence) in the GI tract compared to glucose-fed mice. Interestingly, the effect of FG on intestinal HIF-α stability was similar to fructose alone (Figure S1A-E), suggesting dietary fructose stabilizes intestinal HIF-α compared to glucose. Moreover, we noticed that FG feeding led to a trend of higher blood glucose level (Figure S1F). Similarly, this finding was further validated with a higher dose of sugar (30% fructose or glucose, w/v) administered to ODD-*luc* mice. Notably, high-fructose-induced HIF-α stabilization is more evident in the proximal intestine but is less prominent or has no effect on the distal intestine compared to glucose (Figure S1G-J), likely owing to excess fructose metabolism by gut microbiota,^6^ which leads to decreased HIF-α stability.

^25^ To exclude the confounding factor of varied dietary sugar preferences (fed *ad libitum*) ^26^ on intestinal HIF-α stability, we administered equal amounts of F, G, or FG to ODD-*luc* mice via daily oral gavage for three consecutive days. A previous ^13^C labeled isotope tracer study in mice showed that the majority of labeled fructose metabolite, F1P, accumulated in the SI, peaked 15 minutes after oral gavage of ^13^C-fructose, and disappeared within about 45 minutes. Unlike fructose, the majority of ^13^C labeled glucose 6-phosphate (G6P) accumulated in the muscle, peaked 30 minutes after oral gavage of ^13^C-glucose, and disappeared within about 45 minutes. ^6^ Therefore, we performed BLI of entire GI tract between 15 min to 45 min after oral sugar gavage. We found that daily sugar gavage recapitulates the findings in the chronic feeding model. Notably, glucose gavage resulted in a rapid and robust decrease in luciferase activity in 15 minutes throughout the entire GI tract compared to F or FG mixture (Figure 1A-D). Female mice showed a similar trend of alterations in intestinal HIF-α stability compared to male mice (Figure S1K). Similar findings were also observed in older mice (Figure S1L, M).

**Figure 1.**
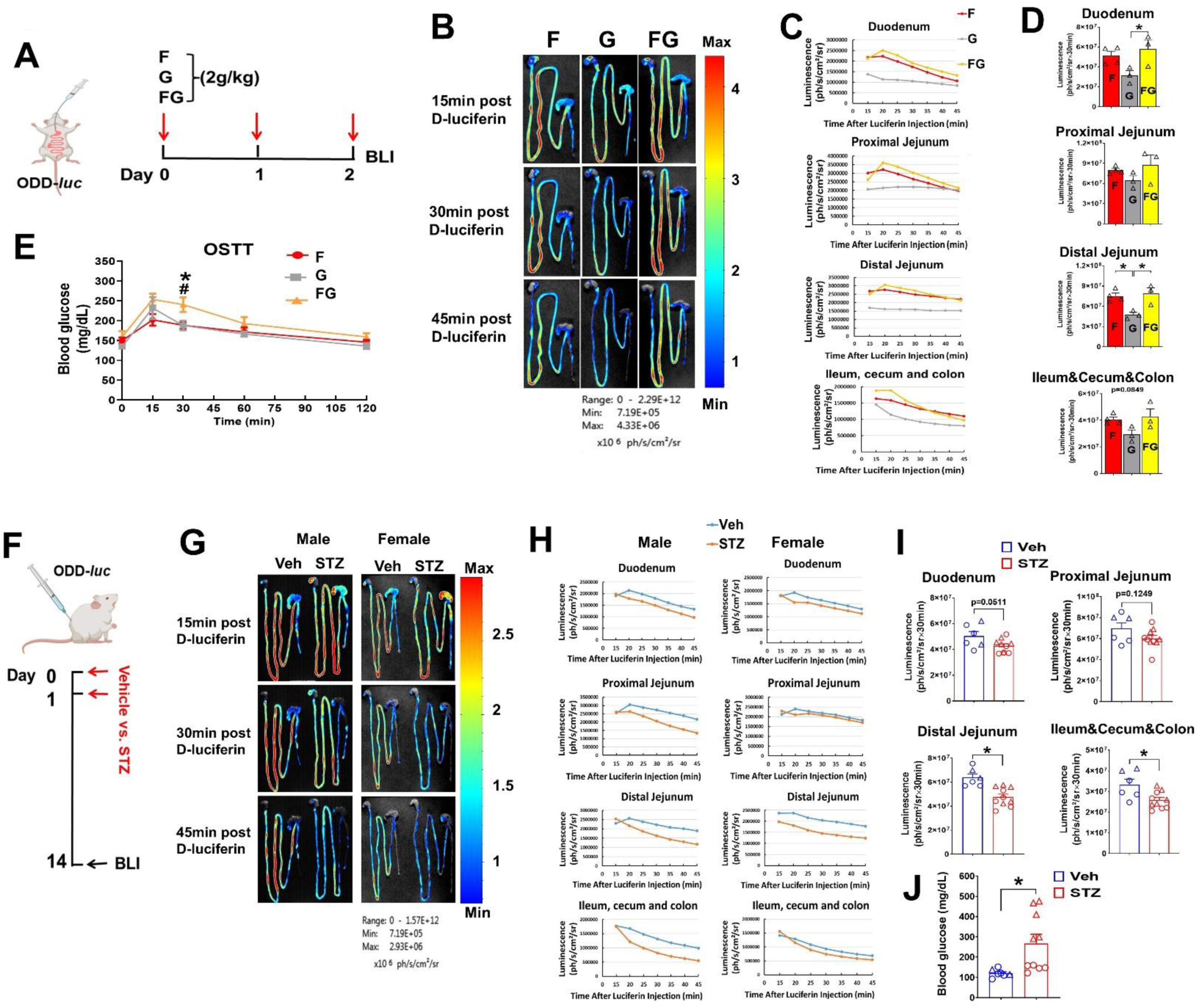
Fructose aberrantly stabilizes intestinal HIF-α. (A) Schematic experimental design for sugar oral gavage on intestinal HIF-α stability. (B) Representative BLI images of HIF-α luciferase activity in the GI tract of 10-12-week-old male ODD-*luc* mice with daily gavage of indicated sugars (at dose of 2 g/kg) for 3 days. (C) Intestinal kinetic curve of HIF-α luciferase activity in terms of luminescence (ph/s/cm²/sr×30min). (D) Calculated area under curve (AUC) of (C) (n=3-4). (E) Oral sugar tolerance test (OSTT) with indicated sugars at the dose of 2g/kg BW in 20% saline solution (n=5-6). **\*** vs F; # vs G. (F) Schematic experimental design for STZ-induced hyperglycemia model. (G) Representative images of GI tract BLI for HIF-α luciferase activity. (H) Intestinal kinetic curve of measured HIF-α luciferase activity in terms of luminescence (ph/s/cm²/sr×30min) (n=2-5). (I) Calculated area under curve (AUC) of (H) (n=6-10). (J) Blood glucose level (n=6-10). Males are designated as triangles, and females are designated as circles (I, J). Data are presented as the mean ± SEM, **\***p<0.05, one-way ANOVA followed by Tukey’s multiple comparison test or unpaired t test. F, fructose; G, glucose; FG, fructose + glucose at a 1:1 mixture; Veh, vehicle; STZ, Streptozotocin.

Hyperglycemia is a common feature induced by dietary sugar intake. We further performed an oral sugar tolerance test (OSTT). It appears that FG results in a higher glycemic effect compared to either fructose or glucose alone (Figure 1E). Given that hyperglycemia is a causal factor of intestinal barrier dysfunction, ^27^ and HIF-1 is a master regulator of intestinal barrier function,^28^ we hypothesized that hyperglycemia inhibits intestinal HIF-α stability. For this purpose, we injected streptozotocin (STZ) intraperitoneally into male adult ODD-*luc* mice to induce hyperglycemia, a commonly used type I diabetes mellitus model. ^27^ As shown in Figure 1F-J, STZ treatment led to robustly elevated blood glucose level, concomitant with significantly reduced HIF-1α luciferase activity in ODD-*luc* mice; this effect was more pronounced in male mice than in female mice. We further validated these results in C57BL/6J mice by PMDZ staining (Figure S1N-P). Despite a higher and sustained hyperglycemic effect of FG compared to either fructose or glucose (Figure 1E), the effect of FG on intestinal HIF-α stability is similar to fructose alone. This suggests fructose preserves HIF-α stability under hyperglycemia, highlighting the specific role of fructose in HIF-α stability.

Fructose can be generated endogenously from glucose via the polyol pathway, catalyzed by aldose reductase (AR), with more than 30% glucose being converted to fructose under hyperglycemia. ^29,30^ Given the role of fructose in HIF-α stabilization under hyperglycemia, we further examined the role of endogenous fructose in the regulation of intestine HIF-α stability. To this end, we pretreated mice with an AR inhibitor, zopolrestat, ^31^ one hour before glucose gavage. AR inhibition did not result in an apparent alteration of intestinal HIF-α stability in the mice with glucose bolus administration (Figure S1Q-T). These data suggest that endogenous fructose is not sufficient to alter intestinal HIF-α stability in glucose-induced hyperglycemia. Moreover, unlike dietary fructose, which is primarily metabolized by the SI, a large portion of circulating fructose is metabolized in the liver and kidney in addition to the SI. ^6^ Thus, it is not surprising that inhibition of endogenous fructose does not significantly affect the intestinal HIF-α stability in the presence of bolus glucose administration.

Collectively, these data suggest that fructose and glucose differentially regulate intestinal HIF-α stability.

### HIF-α stability exhibits regional differences in the intestine

The anatomy and physiology of the intestine exhibit regionalization with each region characterized by distinct functions orchestrated by a regionally specialized immune system, microbiome, and enterocytes. ^32–34^ Physiologic hypoxia is a key feature of a healthy gut, and HIFs (HIF-1α; HIF-2α) serve as important transcriptional regulators in adaptation to low oxygen tensions.^15^ Taking advantage of the HIF-α reporter mice, we were able to visualize physiologic hypoxia across the entire GI tract through *ex vivo* BLI. Notably, we found that HIF-α luciferase activity is unequally distributed along the normal healthy gut, with the highest levels in the jejunum and the lowest in the distal intestine. Furthermore, regionalized HIF-α luciferase activity remained stable across fed and fasting states despite fluctuations in blood glucose and cecum mass (Figure 2A, B, Figure S2A), suggesting that feeding cycles do not alter intestinal physiologic hypoxia under normal physiological conditions. Consistently, the mRNA expression of *Hif-1α* and *Hif-2α* and their target genes are markedly increased in the jejunum compared to the ileum in C57BL/6J mice. In line with nutrient absorption functions, the mRNA expression of fatty acid transport proteins (FATPs) and *Cd36*, which mediate long-chain fatty acid uptake, ^35^ is markedly increased in the jejunum compared to the ileum (Figure 2C).

**Figure 2.**
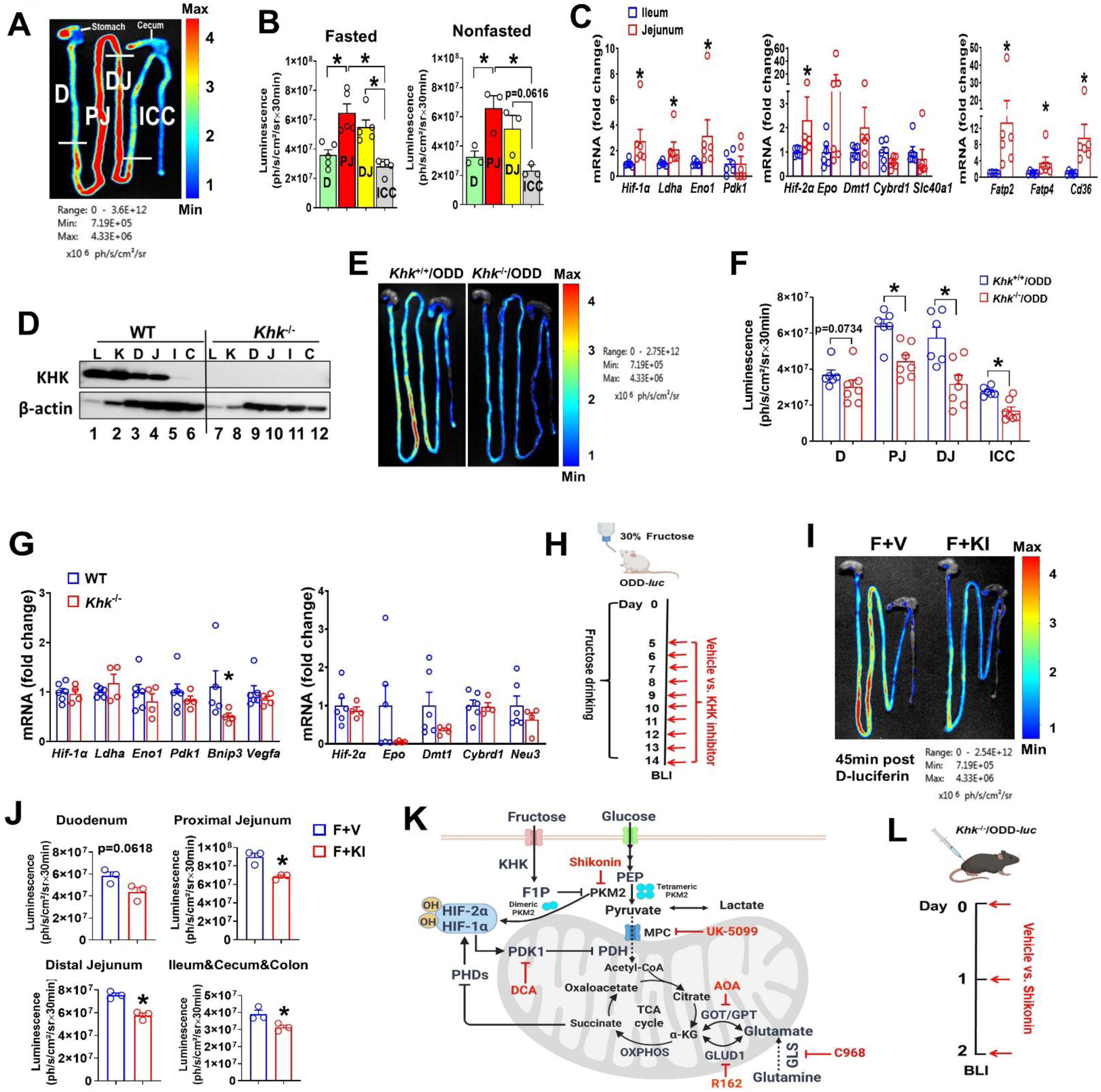
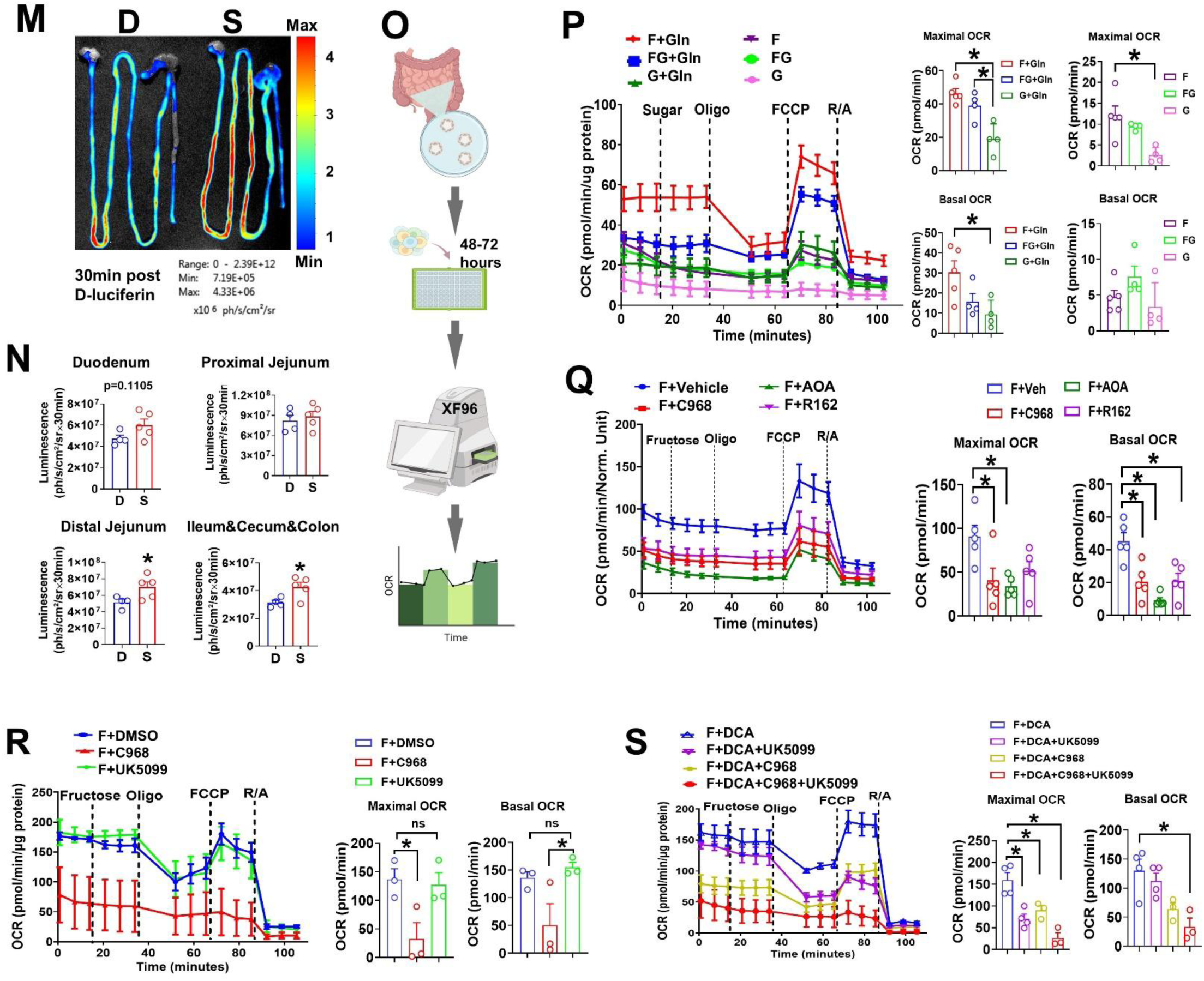
HIF-α stability exhibits regional differences in the intestine and KHK is necessary for maximal steady-state intestinal HIF-α stability. (A) Representative image of *ex vivo* BLI in fasted adult male ODD-*luc* mice. (B) Intestinal HIF-α luciferase activity in fasted and nonfasted normal ODD-*luc* mice(n=3-5). (C) RT-qPCR results of mRNA expression of HIF-1α (left), HIF-2α (middle) target genes, and fatty acid transporters (right) in the jejunum and ileum of male adult C57BL/6J mice (n=5-6). (D) KHK protein expression by western blot. L, liver; K, kidney; D, duodenum; J, jejunum; I, ileum; C, colon. Representative *ex vivo* BLI images of GI tract (E) and quantitative intestinal HIF-α luciferase activity (F) in adult male *Khk^+/+^*/ODD-*luc* and *Khk^-/-^/*ODD-*luc* mice (n=6-7). (G) mRNA expression of HIF-1α (left), HIF-2α (right) target genes in the jejunum of male adult *Khk*^-/-^ mice and WT littermates by RT-qPCR (n=4-6). (H) Schematic experimental design for KHK inhibitor on intestinal HIF-α stability. (I) Representative BLI images of GI tract in 8-week-old male ODD*-luc* mice treated with KHK inhibitor (KI) or vehicle (V) in the presence of 30% (w/v) fructose (F) via drinking water for 2 weeks *ad libitum*. (J) Quantitative intestine HIF-α luciferase activity (n=3). (K) Schematic illustration of fructose induced metabolic reprogramming in mouse SI enterocytes. (L) Schematic experimental design for a PKM2 inhibitor on intestinal HIF-α stability of *Khk^-/-^/*ODD-*luc* mice. (M) Representative BLI images of GI tract in 14-week-old male *Khk^-/-^/*ODD-*luc* mice treated with a selective PKM2 inhibitor, shikonin (S) or DMSO (D) via IP injection. (N) Quantitative intestine HIF-α luciferase activity (n=4-5). (O) Schematic workflow of mitochondrial stress test of isolated SI organoids by Seahorse XF96 analyzer. (P) Oxygen consumption rate (OCR) of proximal SI organoids from male adult wild type mice by Seahorse XF96 analysis with or without 2 mM glutamine in the presence of 150µm sodium pyruvate. (Q) (R) OCR in the absence or (S) presence of DCA with XF DMEM assay medium (sugar-free) containing 2mM sodium pyruvate and 2mM glutamine with sequential injection of fructose 15 mM, oligomycin 5 µM, FCCP 2 µM, rotenone 2 µM, and antimycin A 2 µM (n=3-5 replicates). F, fructose (15mM); FG, fructose: glucose=1:1 (15mM); G, glucose (15mM); MPC, mitochondrial pyruvate carrier; PDH, pyruvate dehydrogenase; PDK1, pyruvate dehydrogenase kinase 1; PEP, phosphoenolpyruvate; GOT, glutamic-oxaloacetic transaminase; GPT, glutamic-pyruvic transaminase; GLUD1, glutamate dehydrogenase 1; GLS, glutaminase; C968, compound 968 (10µM), a glutaminase inhibitor; AOA, Aminooxyacetate (1mM), a pan-transaminases inhibitor; R162, a glutamate dehydrogenase 1 inhibitor (40µM); UK-5099, an MPC inhibitor(10µM); DCA, dichloroacetate (20mM), a PDK1 inhibitor. Data are presented as the mean ± SEM. * p<0.05, one-way ANOVA followed by Tukey’s multiple comparison test or unpaired t test or Mann-Whitney test.

### The fructose metabolizing enzyme, KHK, is necessary for maximal steady-state intestinal HIF-α stability

To better understand the mechanisms by which fructose metabolism increases GI tract HIF-α stabilization, we next generated *Khk*^-/-^ mice in which both isoforms, *Khk*-A and *Khk*-C, were deleted using CRISP/Cas9 (Figure S2B). ^36^ Of note, the KHK enzyme is primarily expressed in the duodenum and jejunum of wild-type (WT) mice but is very weakly expressed in the ileum and undetectable in the colon (Figure 2D), which is positively associated with HIF-α stability. Moreover, KHK expression is consistent with the bulk of dietary fructose being metabolized in the upper GI tract. ^6^ Given the fructose-specific effects on HIF-α stabilization (Figure 1) in the GI tract, coupled with the fact that KHK expression is primarily restricted to the upper GI tract (Figure 2D), we next asked whether loss of KHK specifically reduces HIF-α stability in the intestines of *Khk*^-/-^ mice. To test this, we crossed ODD-*luc* mice with *Khk*^-/-^ mice and their *Khk*^+/+^ littermates,^37,38^ generating *Khk^+/+^*/ODD-*luc* and *Khk^-/-^/*ODD-*luc* mice, respectively. Whole-body BLI confirmed successful ODD-*luc* luciferase expression in both *Khk*^+/+^ and *Khk*^-/-^ mice (Figure S2C). Interestingly, loss of KHK resulted in significantly less HIF-α stability in nearly the entire GI tract; not just the upper portion (duodenum; proximal jejunum) where KHK expression is localized, but also in the distal intestine, which lacks KHK expression (Figure 2E, F). These results suggest that KHK-mediated fructose metabolism can regulate HIF-α stability in a cell non-autonomous manner, likely by increasing microbiota consumption of fructose, resulting in distal HIF-α instability. ^6,28^ Notably, mice in this study were fed only normal mouse chow, which contains small amounts of sucrose, suggesting that the intestinal HIF-α stabilization effect of fructose/KHK does not require excess dietary fructose and can be activated even at low levels by steady-state normal chow consumption. To determine if decreased HIF-α stability in the upper intestine of *Khk*-deficient mice impairs HIF-1α and/or HIF-2α transcription, we harvested intestinal jejunums from *Khk*^+/+^ and *Khk*^-/-^ mice and evaluated the expression of several HIF-1α and HIF-2α target genes by qRT-PCR. While the expressions of *Hif-1a* and *Hif-2a* mRNAs were not affected by *Khk* deficiency, the HIF-1α target gene *Bnip3* and the HIF-2α target genes, erythropoietin (*Epo*) and iron transporter *Dmt1*, were reduced (Figure 2G). We further show that a pharmacologic KHK inhibitor decreases intestinal HIF-α stability *in vivo* during fructose feeding via drinking water (Figure 2H-J). Collectively, these data suggest that the fructose-metabolizing enzyme, KHK, is essential for the maintenance of intestinal HIF-α stabilization and transcriptional activity.

Given that PKM2 transactivates HIF-α, ^21^ and the fructose metabolite F1P inhibits PKM2 activity, ^9^ we hypothesized that fructose-induced HIF-α stability is mediated by PKM2 (Figure 2K). To test this, *Khk^-/-^*/ODD-*luc* mice were treated with a selective PKM2 inhibitor, shikonin. ^39,40^ Intestinal HIF-α luciferase activity was significantly increased by shikonin compared to vehicle-treated mice (Figure 2L-N) (i.e., it rescues the reduced HIF-α stability as shown in Figure 2E). HIF activation dictates glutamine entry into the tricarboxylic acid (TCA) cycle for anaplerosis via pyruvate dehydrogenase kinase 1 (PDK1)-mediated inhibition of pyruvate dehydrogenase (PDH). ^41,42^ This effect is likely enhanced by F1P-induced inhibition of PKM2 catalytic activity, which leads to reduced pyruvate production. Consequently, this metabolic reprogramming may cause glutamine-derived succinate accumulation,^9^ which competitively inhibits α-ketoglutarate (α-KG)-dependent prolyl hydroxylase (PHD) activity, thereby preventing HIF-α degradation and increasing HIF-1α stability. ^43,44^ To further understand the metabolic effects of fructose on HIF-α stabilization, we performed a mitochondrial stress test using isolated crypt organoids from the SI of mice (Figure 2O-S). The mitochondrial stress test showed that fructose markedly increases mitochondrial respiration compared to glucose as shown by the oxygen consumption rate (OCR) in the presence of glutamine (Figure 2P). This effect was inhibited by inhibitors of the glutaminolysis pathway (compound 968, AOA and R162) (Figure 2Q) but not by the mitochondrial pyruvate carrier (MPC) inhibitor, UK-5099 (Figure 2R), suggesting that fructose induced oxidative phosphorylation is glutamine-dependent and pyruvate-independent. To test whether fructose-induced metabolic reprogramming is mediated by PDK1, SI organoids were pretreated with dichloroacetate (DCA), a PDK1 inhibitor, which activates PDH and catalyzes the conversion of pyruvate to acetyl-CoA, allowing pyruvate to enter the TCA cycle. ^45^ In the presence of DCA, fructose-induced mitochondrial respiration (OCR) was inhibited by both compound 968 and UK-5099, showing a great extent of inhibition through the combinatorial effects of compound 968 and UK-5099 (Figure 2S). This supports our hypothesis that fructose aberrantly stabilizes HIF and that this prevents pyruvate from entering the TCA cycle via PDK1-mediated PDH inactivation (Figure 2K). Given that fructose metabolism is dictated by HIF-1α through SF3B1-mediated alternative splicing to KHK-C,^12^ our study further demonstrates that fructose metabolism stabilizes HIF-1α. Because KHK is required for this stabilization; this suggests a feedforward loop in which the KHK-HIF axis sustains fructose metabolism under hypoxia.

### KHK deficiency prevents fructose-induced increases in plasma iron levels while disrupting systemic iron homeostasis

Given our findings that fructose/KHK promotes intestinal HIF-α stabilization and intestinal HIF-2α are important determinants of intestinal iron absorption,^16,17^ we next asked whether dietary fructose influences iron homeostasis and whether KHK is necessary. As shown in Figure 3A, B, dietary fructose consumption significantly increases plasma iron levels in WT mice, and this effect is completely abrogated in *Khk*^-/-^ mice. Conversely, *Khk*^-/-^ mice exhibit significant increases in plasma EPO levels compared to WT controls (Figure 3C). Concomitantly, fructose consumption preserves intestinal physiological hypoxia in WT mice, whereas it fails to do so in *Khk*^-/-^ mice, as shown by duodenum PMDZ IHC staining (Figure S3A). Moreover, plasma iron levels are significantly lower in *Khk*^-/-^ mice compared to WT littermates on an iron-sufficient diet across different age groups (4 months and 14 months) (Figure 3D). Importantly, and very consistent with our observed requirement for fructose/KHK in driving steady-state HIF-2α-dependent *Epo* and *Dmt1* expression (Figure 2G), *Khk*^-/-^ mice exhibit spontaneous mild hypochromic anemia (Figure 3E, F) and downregulated hepatic *Hamp* gene expression, as shown by both RT-PCR and RNA-seq (Figure 3G), suggesting disrupted systemic iron homeostasis. Notably, markedly reduced neutrophil numbers and percentages are found in the blood of *Khk*^-/-^ mice, while total WBC numbers remain largely unchanged (Figure 3H). Bulk RNA-seq analysis identified that pathways involving the ribosome and oxidative phosphorylation in the jejunum, ileum and liver, as well as metabolism of xenobiotics by cytochrome P450, drug metabolism-cytochrome P450, and ribosome biogenesis in eukaryotes in the duodenum, are among the top ten downregulated KEGG pathways in *Khk*^-/-^ mice compared to WT mice (Figure 3I). All of these pathways require Fe-S clusters or heme iron as cofactors. ^46^ Furthermore, GO terms related to DNA synthesis and mitochondrial respiration are among the top ten downregulated GO terms in *Khk*^-/-^ mice (Figure 3J), which also require iron. Collectively, these RNA-seq data support the systemic effect of iron deficiency in inducing transcriptomic changes.

**Figure 3.**
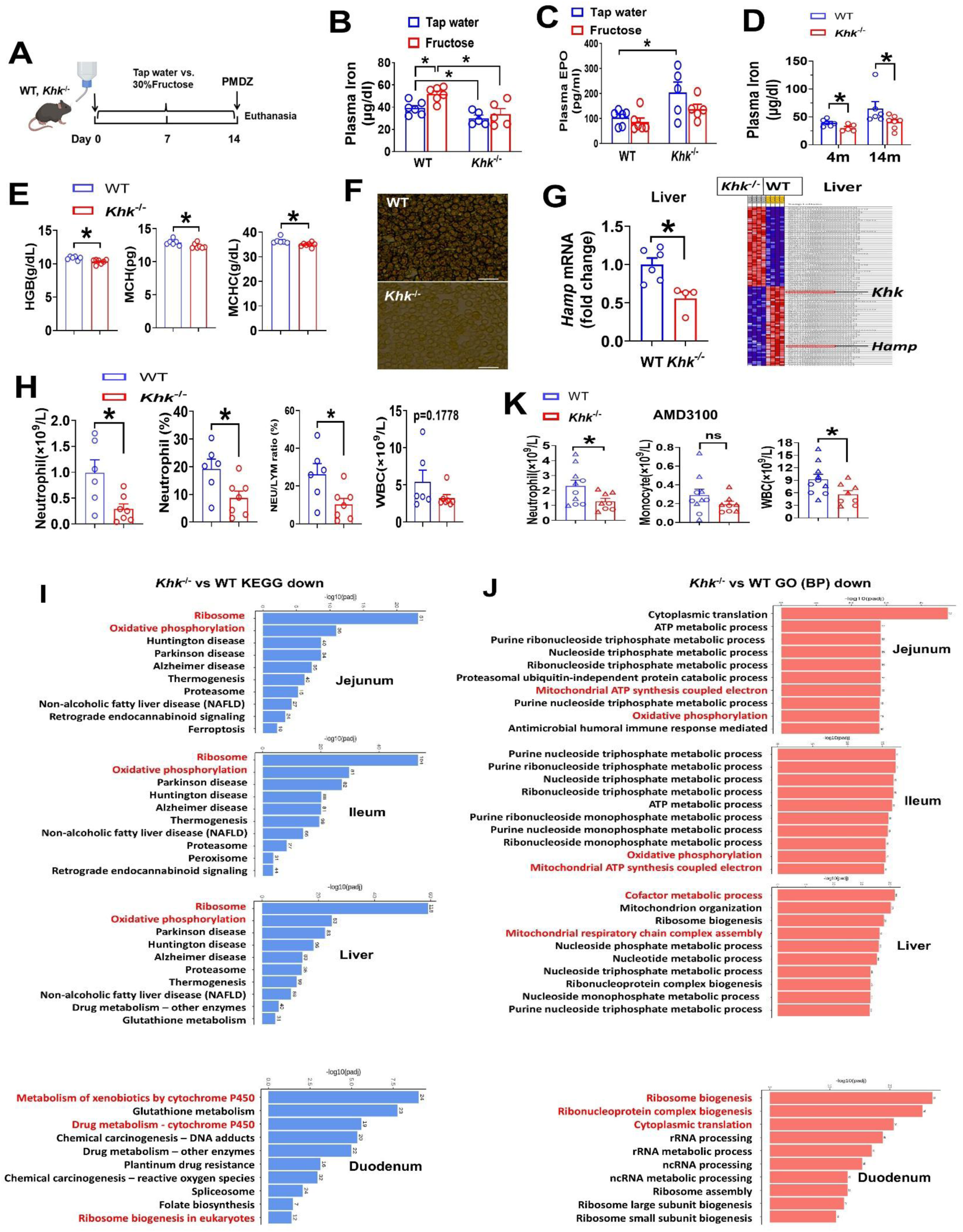
KHK deficiency prevents fructose-induced increases in plasma iron levels while disrupting systemic iron homeostasis. (A) Schematic experimental design for fructose feeding on iron levels. (B) Plasma iron and (C) plasma EPO levels in 4-month-old WT and *Khk*^-/-^ mice (n=5-6). (D) Plasma iron in young/aged male mice (n=5-7). Plasma iron level of 4-month-old WT and *Khk*^-/-^ mice in (D) shared with mice with tap water in (B). (E) Blood HGB, MCH, and MCHC levels of 14-month-old mice by complete blood count (CBC) analysis (n=5-7). (F) Representative images of blood smear. Scale bar, 40µm. (G) The expression of hepatic *Hamp* mRNA determined by RT-qPCR (left) (n=4-6) and heatmap of differential gene expression by bulk RNA-seq (right) (n=4). Left 4 column, *Khk*^-/-^ mice; right 4 column, WT mice. Expression values are represented as colors, where the range of colors (red, pink, light blue, dark blue) shows the range of expression values (high, moderate, low, lowest), which represent the significant level of enrichment. (H) Blood neutrophil numbers and percentage of 14-month-old mice by CBC analysis (n=5-7). (I) Top ten downregulated KEGG pathways and (J) GO terms by bulk RNA-seq analysis (n=4). (K) Blood neutrophils, monocytes and WBCs after 2 hours AMD3100 injection (n=8-10). Data are presented as the mean ± SEM. *p<0.05, two-way ANOVA followed by Tukey’s multiple comparison test or unpaired t-test or Mann-Whitney test. EPO, erythropoietin; HGB, hemoglobin; MCH, mean corpuscular hemoglobin; MCHC, mean corpuscular hemoglobin concentration; WBC, total white blood cell count.

Because KHK deficiency results in decreased plasma iron levels and blood neutrophils, and given the critical role of plasma iron in neutropoiesis, ^47,48^ we hypothesized that KHK deficiency-induced iron deficiency might inhibit hematopoiesis. Thus, we further analyzed bone marrow neutrophils; however, there was no significant difference between WT and *Khk*^-/-^ mice in the percentage of CD45^+^CD11B^+^Ly6G^+^ bone marrow cells, as determined by flow cytometry (Figure S3B-D). Since HIF-1α is required for hematopoietic stem cell (HSC) mobilization,^49^ and KHK is necessary for HIF-α stabilization, we investigated whether KHK-deficiency-induced neutropenia results from reduced HSC mobilization from the bone marrow. In response to a CXCR4 antagonist AMD3100, ^50^ blood neutrophils were significantly reduced in *Khk*^-/-^ mice compared to WT mice, suggesting *Khk* deficiency inhibits HSC mobilization, presumably via the KHK/HIF axis (Figure 3K).

### KHK deficiency inhibits iron absorption in a HIF-2α-dependent manner

HIF-2α signaling is required for iron absorption. ^16^ Both a low iron diet ^51^ and hemolytic anemia ^18^ activate intestinal HIF-2α, leading to hyperabsorption of iron. To address whether KHK deficiency results in reduced iron absorption in a HIF-2α-dependent manner, 4-6-week-old male and female *Khk*^-/-^ mice and WT littermates were placed on a low-iron diet or iron-replete diet for 3 weeks (Figure 4A). ^18,23^ IHC staining showed that duodenal HIF-2α was markedly activated by a low-iron diet in WT mice, and that effect was abrogated in *Khk*^-/-^ mice (Figure 4B). Consistently, duodenal HIF-2α target gene expression exhibited a trend of upregulation on a low-iron diet in WT mice, which was blunted in *Khk*^-/-^ mice (Figure S4A). In line with this, plasma hepcidin levels trended toward an increase on a low-iron diet in WT mice compared to those on an iron-replete diet, yet this difference did not reach statistical significance (Figure S4B). On the contrary, plasma iron levels were reduced in *Khk*^-/-^mice fed on an iron-replete diet compared to WT littermates. However, levels did not differ between *Khk*^-/-^ mice and WT littermates when exposed to a low-iron diet (Figure 4C), suggesting that KHK deficiency inhibits iron absorption.

**Figure 4.**
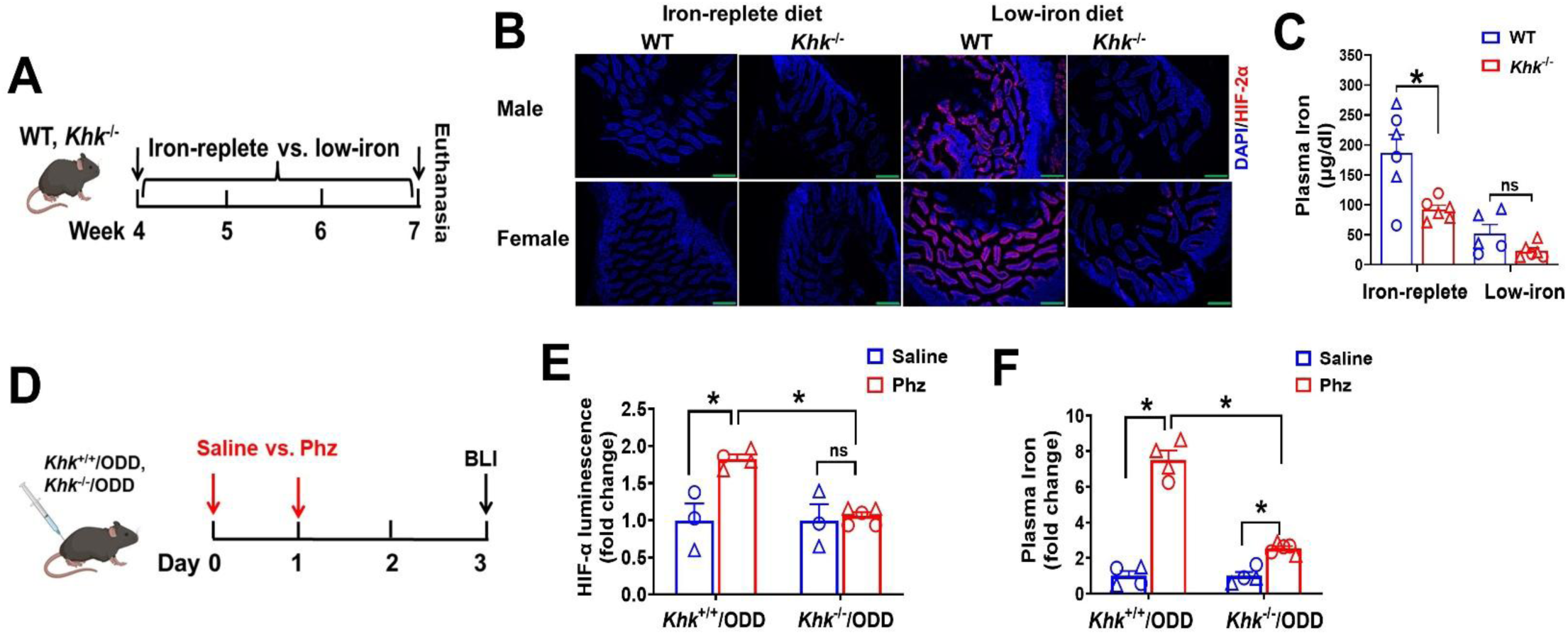
KHK deficiency inhibits iron absorption in a HIF-2α-dependent manner. (A) Schematic experimental design for low-iron diet-induced iron deficiency model. (B) Representative images of duodenal HIF-2α immunofluorescence (IHC) staining (n=5-6). Scale bar, 200µm. (C) Plasma iron levels (n=5-7). (D) Schematic experimental design for Phz-induced hemolytic anemia model. (E) Duodenal HIF-α luciferase activity (n=3-5). (F) Alterations of plasma iron in response to Phz (n=4-5). Data are presented as the mean ± SEM. * p<0.05, two-way ANOVA followed by Tukey’s multiple comparison test. Males are designated as triangles, and females are designated as circles (C, E, F). Phz, phenylhydrazine.

To further validate this, *Khk^+/+^*/ODD-*luc* and *Khk^-/-^/*ODD-*luc* mice were subjected to a Phz-induced hemolytic anemia model (Figure 4D, Figure S4C). Duodenal HIF-α luciferase activities were significantly increased in Phz-treated *Khk^+/+^*/ODD-*luc* mice compared to vehicle-treated mice, which was abrogated in *Khk^-/-^/*ODD-*luc* mice (Figure 4E). Phz treatment led to a 7-fold increase in plasma iron level in *Khk^+/+^*/ODD-*luc* mice compared to vehicle-treated mice, but to a much lesser extent in *Khk^-/-^/*ODD-*luc* mice (∼2-fold increase) (Figure 4F). In addition, *Khk^-/-^*ODD-*luc* mice exhibited reduced splenic iron compared to *Khk^+/+^*/ODD-*luc* mice with saline treatment, as shown by Perls’ Prussian blue iron staining; however, the difference was not evident in Phz-treated mice (Figure S4D). Collectively, these data support the concept that KHK deficiency inhibits iron absorption in a HIF-2α-dependent manner.

### The KHK/HIF axis plays a critical role in MASH/HCC progression

To determine the role of the KHK-HIF axis in MASH and HCC progression, *Khk^+/+^*/ODD-*luc* and *Khk^-/-^*/ODD-*luc* mice were subjected to a murine MASH/HCC model induced by neonatal STZ injection and a high-fat diet (STAM mouse model), which combines diabetes and hepatic steatohepatitis and recapitulates the major genetic and phenotypic features of human MASH. ^52,53^ Given that KHK deficiency results in decreased intestinal HIF-α activity in normal physiological condition (Figure 2E-G), we further tested whether KHK-deficiency inhibits intestinal HIF-α stability in a pathophysiological MASH/HCC STAM model. ^52^ Indeed, intestinal HIF-α stabilization was significantly reduced in male *Khk^-/-^*/ODD-*luc* mice compared with *Khk^+/+^*/ODD-*luc* mice at 8 weeks of the STAM model, and this was associated with attenuated hepatic steatosis (Figure 5A-D, Figure S5A, B) as well as mitigated tumorigenesis (Figure 5E-G, Figure S5C, D). Consistent with this, male *Khk^-/-^* mice exhibited attenuated tumorigenesis in the DEN/HCC model (Figure 5H-K, Figure S5E, F).

**Figure 5.**
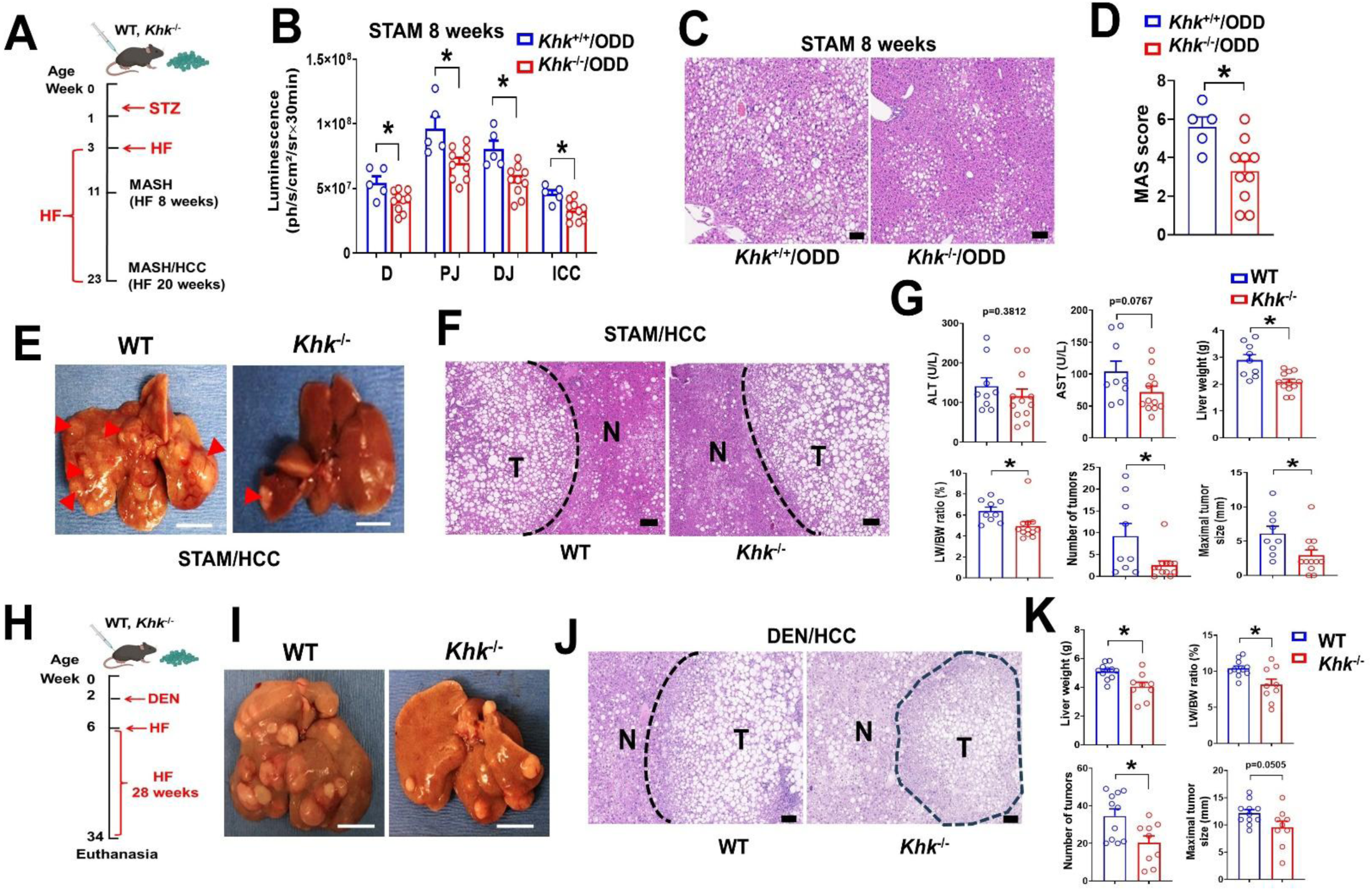
The KHK/HIF axis plays a critical role in MASH/HCC progression. (A) Schematic timeline of STAM MASH/HCC model. (B-D) Male *Khk^+/+^*/ODD-*luc* and *Khk^-/-^/*ODD-*luc* mice were subjected to STAM model for 8 weeks. (B) Intestine HIF-α luciferase activity by BLI at 8 weeks (n=5-10). (C) Representative images of liver H&E staining. (D) MASLD activity score (MAS). Representative gross morphology of STAM/HCC mice liver (E), representative images of liver H&E staining (F) and HCC development (G) in male WT and *Khk*^-/-^mice subjected to STAM/HCC model for 20 weeks (n=9-12). (H) Schematic timeline of DEN/HCC model. Representative gross morphology of liver (I) and Liver H&E staining (J) and HCC development (K) in male WT and *Khk*^-/-^mice subjected to DEN/HCC model (n=9-11). Data are presented as the mean ± SEM. * p<0.05, unpaired t test. Scale bar, 100µm (C, F, J); Red arrow heads in (E) denotes tumor. Scale bar, 1cm (E, I).

We further compared intestinal HIF-α luciferase activity between male and female mice in the STAM model. Intestinal HIF-α luciferase activity was higher in male mice (Figure S5G), which was associated with higher liver weights and liver/body weight ratios compared to female mice at 8 weeks (Figure S5H). Moreover, male mice were more prone to developing early HCC compared to females (male, 3/8 vs. female, 0/8), albeit the difference did not reach statistical significance (Figure S5I). Collectively, these data underscore a critical role of the intestinal KHK/HIF axis in promoting MASH/HCC progression.

### Fructose promotes MASH/HCC progression in an iron-dependent manner

Given the differential roles of dietary fructose and glucose in intestinal HIF-α stability and that fructose promotes iron absorption, we further tested whether dietary fructose promotes MASH and HCC progression in an iron-dependent manner. Both male and female WT mice are subjected to the STAM HCC murine model with fructose or glucose supplementation via drinking water. The majority of male STAM mice died within several days of high fructose or glucose feeding, likely owing to hyperglycemia and ketoacidosis. Female STAM mice developed early-stage HCC by the end of the experiment. Dietary fructose feeding promoted MASH/HCC development as shown by significantly increased tumor numbers compared to the glucose-fed mice, which was significantly attenuated with an iron chelator (DFP) (Figure 6A-C, Figure S6A, B). Notably, plasma iron levels were significantly increased by fructose compared to glucose; this was associated with elevated blood HGB, HCT, and RBC levels, which were markedly decreased by DFP (Figure 6D, E). However, the numbers of blood neutrophils, monocytes, WBC and PLT were not significantly changed (Figure S6C). This suggests that fructose specifically impacts erythropoiesis rather than leukocytes or thrombocytes, highlighting the role of fructose in iron homeostasis.

**Figure 6.**
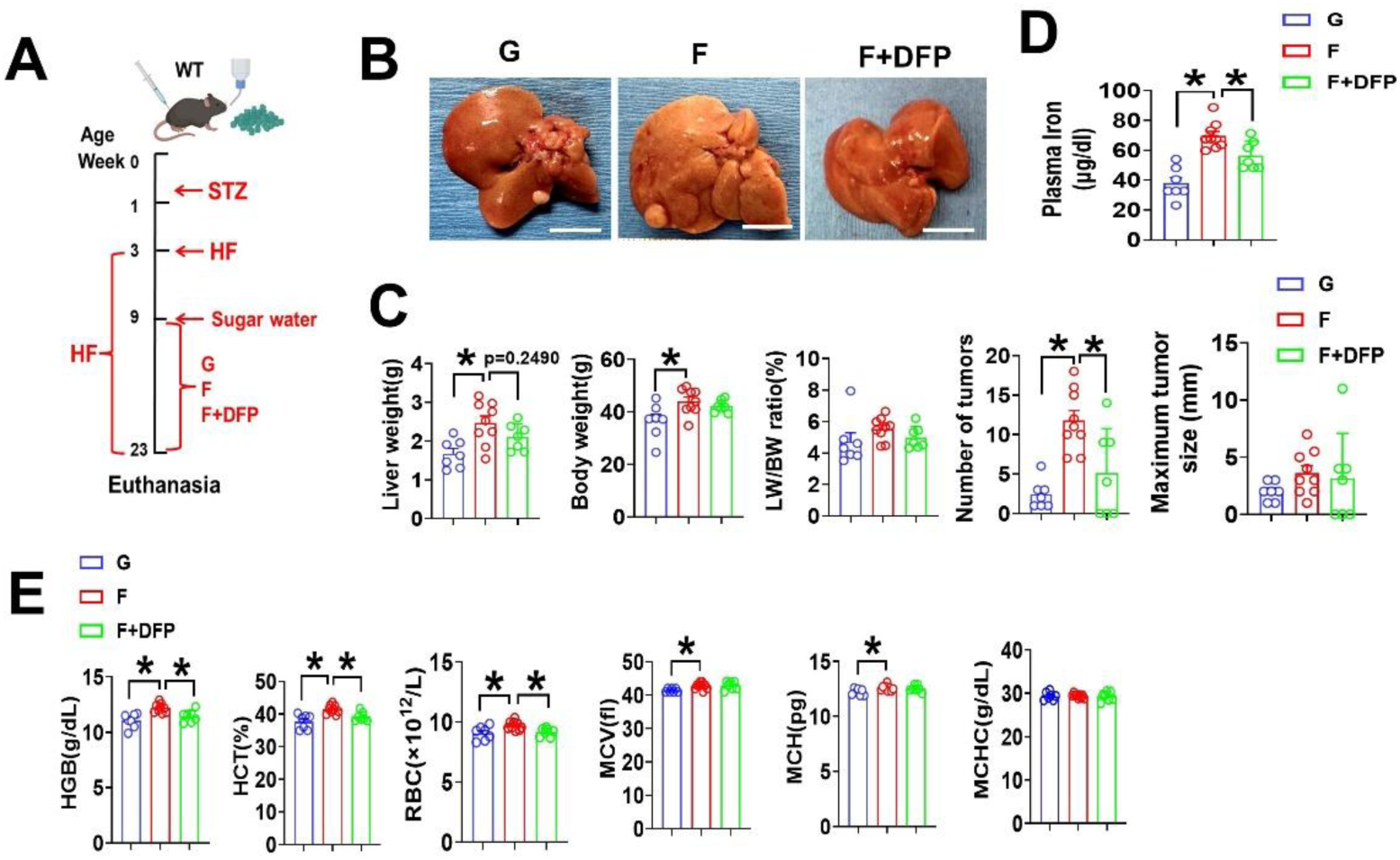
Fructose promotes MASH/HCC progression in an iron-dependent manner. (A) Schematic timeline of STAM MASH/HCC model with sugar supplementation. (B) Representative liver morphology and (C) HCC development in female STAM MASH/HCC model. (D) Plasma iron levels. (E) Blood HGB, HCT, RBC, MCV, MCH and MCHC related to (A-D) (n=7-9). Data are presented as the mean ± SEM. * p<0.05, one-way ANOVA followed by Tukey’s multiple comparison test or unpaired t test. G, glucose; F, fructose; DFP, deferiprone; HF, high-fat diet; STZ, streptozotocin; HGB, hemoglobin; RBC, red blood cell; HCT, hematocrit; MCV, mean corpuscular volume; MCH, mean corpuscular hemoglobin; MCHC, mean corpuscular hemoglobin concentration. All cartoons were created in Biorender.com.

Given that *Dmt1* is a HIF-2α target gene required for intestinal iron absorption, ^17^ we further validated our findings using inducible intestine-specific *Dmt1* knockout mice (*Dmt1*^ΔIEC^) generated by crossing *Dmt1* flox mice with *Villin-Cre-ERT2* mice. Male *Dmt1*^fl/fl^ and *Dmt1*^ΔIEC^ mice were subjected to the DEN/HF HCC model with an additional 15% fructose (w/v) via drinking water (Figure S6D-F). Despite severe anemia induced by intestine-specific *Dmt1* deficiency (Figure S6G-I), the liver tumor number showed a downward trend compared to flox control mice, and the maximal tumor size was not significantly different (Figure S6J, K). Unlike mild anemia induced by *Khk* deficiency or DFP, *Dmt1* deficiency induces severe anemia but proved less protective in MASH and HCC progression.

## DISCUSSION

The distinct metabolic effects of glucose and fructose have been noted. ^4,5,54,55^ A major difference is that fructose is preferentially metabolized under hypoxia. ^9,12,13^ Nonetheless, the underlying mechanisms and their biological significance remain poorly understood. In this study, we demonstrated that fructose metabolism in the SI stabilizes HIF-α via PKM2, a process requiring KHK, which represents a HIF regulatory mechanism. Combined with the finding that HIF-1α regulates KHK splicing via SF3B1,^12^ a complete regulatory loop that dictates and sustains fructose metabolism under hypoxia is established. Moreover, we identified a biological role of the KHK-HIF-2α axis in facilitating iron absorption, which contributes, at least partially, to MASH/HCC progression. Thus, our study revealed a unique fructose sensing pathway through the KHK-HIF-2α axis. These findings not only advance our understanding of the metabolic effects of fructose but also identify a potential therapeutic target for MASH-related HCC.

In adaptation to hypoxia, HIF-1 suppresses mitochondrial respiration and TCA cycle via its target gene PDK1, which limits pyruvate entry into the TCA cycle by inactivating PDH via phosphorylation. ^41,56^ This leads to glutamine, the most abundant circulating amino acid in the body, replenishing the TCA cycle through anaplerosis. ^57^ Indeed, our study demonstrated that fructose promotes glutamine-dependent and pyruvate-independent mitochondrial respiration in SI organoids. This effect is reversed by a PDK1 inhibitor, suggesting that the metabolic effect of fructose activates HIF-1α, leading to glutamine fueling the TCA cycle. In supporting this, Taylor and colleagues ^9^ showed that ^13^C labeled fructose does not enter the TCA cycle in cultured enterocytes under hypoxia with fructose, but instead, glutamine-derived carbon is incorporated into α-KG and succinate, suggesting fructose and/or hypoxia drive glutamine into TCA cycle. They further showed that enterocytes cultured under hypoxia with a fructose-containing medium exhibit robust metabolic activity compared to a glucose medium, as shown by remarkably increased intermediate and fatty acid synthesis, highlighting the critical role of fructose in driving cell metabolism under hypoxia. Moreover, fructose-treated enterocytes result in increased succinate and decreased α-KG compared to glucose, ^9^ which could stabilize HIF through inhibition of PHD-mediated HIF degradation via competitive inhibition of α-KG, a cofactor required for PHD activity. ^43,44^ Similarly, Jones’ study showed that fructose facilitates the incorporation of glutamine-derived carbon into 4-carbon TCA intermediates (fumarate, malate, and aspartate) in macrophages, ^58^ indicating a cell-type specific effect of glutamine metabolism. Therefore, it is likely that fructose metabolism inhibits PKM2 via F1P, leading to HIF-1α transactivation. This prevents pyruvate entry into TCA cycle through PDK1-induced inhibition of PDH, which, in turn, directs glutamine into the TCA cycle. The metabolic fate of glutamine further stabilizes HIF through the accumulation of metabolites such as succinate ^43,44^ or fumarate. ^59^ Furthermore, fructose-induced, glutamine-dependent oxidative phosphorylation consumes more oxygen than glucose, which contributes to HIF stabilization.

Given the metabolic effect of fructose in stabilizing HIF-α, and that *Khk* deficiency reduces HIF-α stability as well as HIF-α transcription—with a more pronounced effect on HIF-2α—we discovered the critical role of fructose/KHK in the regulation of iron homeostasis. Fructose significantly increases plasma iron levels compared to glucose, likely due to enhanced HIF-2α-dependent iron absorption, and it promotes MASH/HCC progression in an iron-dependent manner. Interestingly, we noticed that *Khk* deficiency induces only mild or moderate iron deficiency and anemia, which is different from the severe iron deficiency induced by *Dmt1* deficiency. The precise mechanism underlying how fructose/KHK/HIF-2α fine-tunes iron homeostasis remains elusive, but it probably involves epigenetic or post-transcriptional regulation of HIF-2α target gene. Moreover, *Khk* deficiency, as well as DFP-induced mild iron deficiency, is beneficial in the prevention of MASH/HCC progression. However, this effect was less evident in *Dmt1*^ΔIEC^ mice, likely due to severe iron deficiency caused by *Dmt1* deficiency-induced oxidative stress and mitochondrial dysfunction. ^60^ The protective mechanism of DFP has been attributed to mitophagy. ^61,62^ In supporting this, a study by Softic et al. showed that a high-fat diet plus fructose impairs hepatic mitophagy.^5^

Another interesting finding is that HIF-α stability exhibits regional differences in the intestine coupled with KHK expression. It is well documented that HIF is critical in the maintenance of the gut barrier function in the colon. ^28,63,64^ However, the proximal intestine is colonized with much less abundance of gut microbiota compared to the colon. In contrast, HIF-α stability is higher in the jejunum compared to the colon. This paradoxical phenomenon challenges the paradigm that HIF activity is adapted only for gut barrier function in the jejunum. Given that the jejunum is the major site of nutrient absorption, it is likely that the higher HIF stability in the jejunum is adapted to fulfill the function of nutrient absorption. In line with this, a recent study showed that fructose facilitates lipid absorption. ^9^ From an evolutionary perspective, fructose metabolism was considered an adaptation for stimulating fat storage as a protection mechanism against starvation. ^65^ Given the essential role of iron, our study reveals a more fundamental role of fructose/KHK in the maintenance of iron storage.

In summary, our study identifies a distinct role of dietary fructose in facilitating iron absorption via the KHK-HIF-2α axis, which is an important mechanism underlying the metabolic effects of fructose in the promotion of MASH/HCC progression. Moreover, our study provides preclinical evidence for iron chelators as a potential therapeutic approach. In fact, iron chelators, either alone or in combination therapy with receptor tyrosine kinase inhibitors or chemotherapy, have been shown to be a promising therapeutic strategy for a variety of cancers. ^66,67^

## Limitations of the study

KHK has two isoforms, KHK-A and KHK-C, with KHK-C being the primary isoform for fructose metabolism. ^36,68^ The *Khk*^-/-^ mice used in this study are deficient in both KHK-A and KHK-C. Recent studies suggest KHK-A acts as a protein kinase through phosphorylation of specific substrates involved in tumor metastasis.^69,70^ Given that HIF stability can be regulated by protein kinase, glycogen synthase kinase-3β (GSK3β), leading to HIF-α subunit phosphorylation, subsequent ubiquitination, and degradation, which is independent of oxygen/PHD/VHL, ^71^ further studies to test whether KHK-A directly phosphorylates HIF-α or acts via an intermediate protein to regulate HIF stability are needed.

In addition, BLI-detected HIF stability encompasses the entire intestine wall, including the epithelium, lamina propria, muscularis mucosae, submucosa, and muscularis. Although intestinal epithelial cells are the primary site of fructose metabolism, other cell types, including immune cells, may also metabolize it. ^72^ Thus, the cell-type specific roles and crosstalk between them in the regulation of HIF stability and iron absorption remain elusive.

## RESOURCE AVAILABILITY

### Lead contact

Further information and requests for resources and reagents should be directed to and will be fulfilled by the lead contact upon reasonable request, Ming Song.

### Materials availability

This study did not generate new unique reagents. Mouse lines generated in this study are available upon reasonable request to the Lead Contact with a completed Materials Transfer Agreement.

### Data and code availability

- RNA-seq data have been deposited in the National Center for Biotechnology Information (NCBI) GEO (GSE331146).
- This paper does not report original code.
- Raw data reported in this paper have been deposited on Mendeley Data and DOI was provided on the key resource table.
- Any additional information required to reanalyze the data reported in this paper is available from the lead contact upon reasonable request.

## ACKNOWLEDGMENTS

We thank Drs. Huaiyu Zheng and Chin Ng at the University of Louisville Molecular Imaging Core, as well as Dushan T. Ghooray, and Guangzhong Xu, for their technical support. This study was supported by NIH grants 1R21CA 290420 (M.S., R.A.M.), P20GM113226 (C.J.M., M.S.), and R01CA279748 (R.A.M.); a Jewish Heritage Fund for Excellence Pilot Grant Program at the University of Louisville School of Medicine (M.S.); NIH grants R01HL168198 (B.G.H.), R01HL147844 (B.G.H.), U01AA026934, U01AA026936, and P50AA024337 (C.J.M.); the American Heart Association [23TPA1141824 (B.G.H.)]; and the Veterans Administration (I01BX002996) (C.J.M.). The content is solely the responsibility of the authors and does not necessarily represent the official views of the National Institutes of Health. We thank Marion McClain for editorial assistance.

## AUTHOR CONTRIBUTIONS

Conceptualization, R.A.M., M.S.; data curation, M.S.; formal Analysis, M.S.; investigation, M.S., M.X., M.S.T., B.G.H., R.A.M.; methodology, E.H., M.S.T., B.G.H., M.X.; writing – original draft, M.S.; writing – review & editing, R.A.M. and M.S.; funding acquisition, C.J.M., R.A.M., and M.S.; resources, E.H.; supervision, M.S., R.A.M.

## DECLARATION OF GENERATIVE AI AND AI-ASSISTED TECHNOLOGIES IN THE WRITING PROCESS

The authors declare no generative AI and AI-assisted technologies in the writing process.

## SUPPLEMENTAL INFORMATION

**Document S1. Figures S1–S6, and Tables S1**

## STAR★METHODS

### KEY RESOURCES TABLE

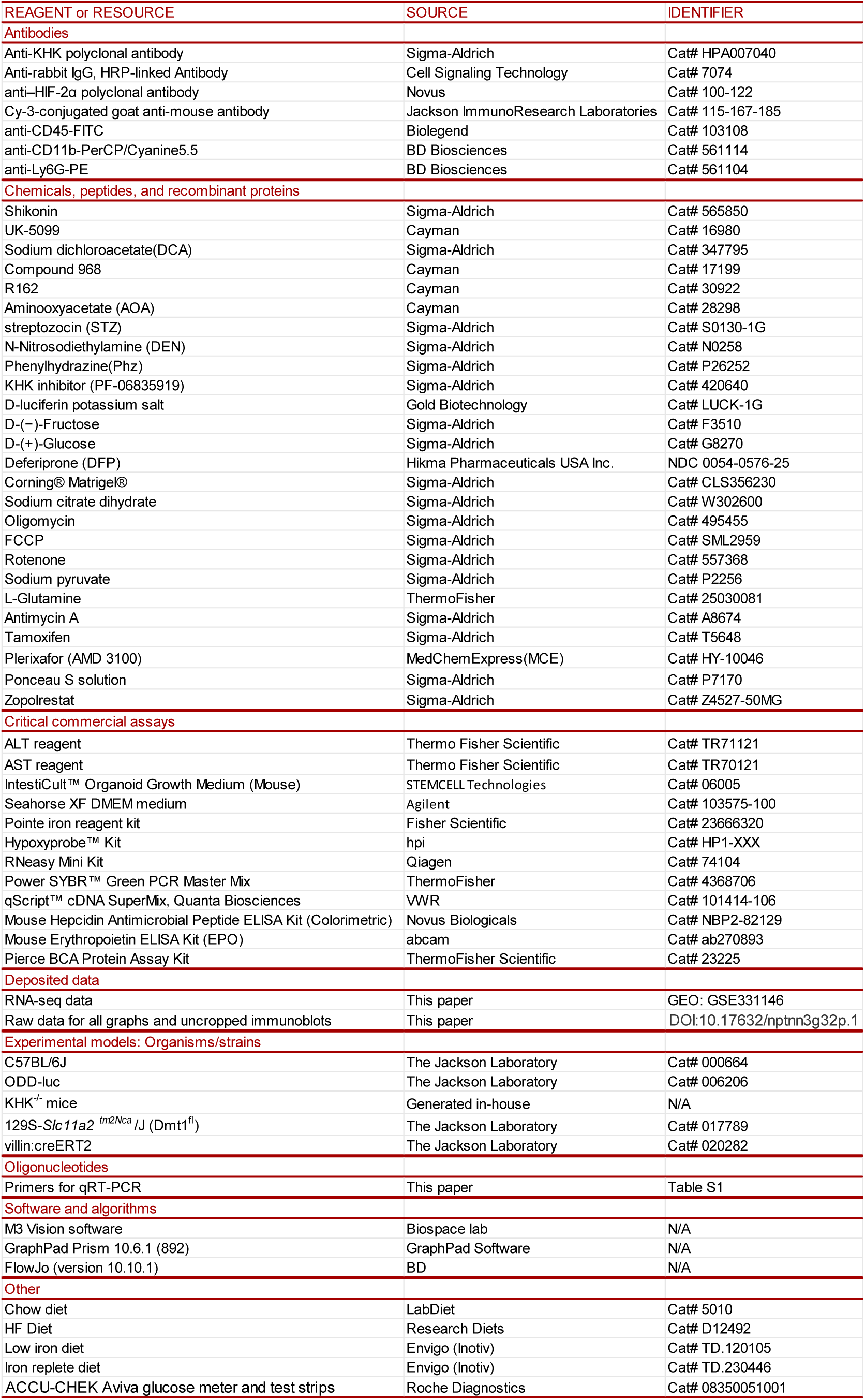

### EXPERIMENTAL MODEL AND STUDY PARTICIPANT DETAILS

#### Mouse strains and husbandry

C57BL/6J, ODD-*luc*, ^22^ *Dmt1* flox^17^ and villin-cre-ERT2 mice ^73^ were obtained from Jackson Laboratory (Bar Harbor, ME). KHK-A/C knockout mice (*Khk*^-/-^) were generated in-house and detailed method was described in METHOD DETAILS. The animals were group-housed in a temperature- and humidity-controlled room with a 12:12 h light-dark cycle and were on a standard pelleted rodent chow diet (5010, LabDiet). Sugar enriched drinking water was changed twice a week. Body weight and food consumption were monitored weekly. At the end of experiment, all the animals were sacrificed under anesthesia with ketamine/xylazine (100/10mg/kg I.P. injection) after fasting for 4 hours. Blood was collected from the inferior vena cava, and citrated plasma was stored at -80°C for further analysis. Portions of liver and intestine tissue were fixed with 10% formalin, and the other tissues were snap-frozen with liquid nitrogen. All animal studies were approved by the University of Louisville Institutional Animal Care and Use Committee, which is certified by the American Association of Accreditation of Laboratory Animal Care.

### METHOD DETAILS

#### Generation of *Khk*-A/C knockout mice

Guide RNA design and CRISPR/Cas9 Assembly: The CRISPR/Cas9 system was used to generate a global knockout of both isoforms A and C of Ketohexokinase (Khk). Three single guide RNA’s (sgRNA) were designed using crispor.tefor.net and the Custom Alt-R CRISPR-Cas9 guide RNA tool from IDT and were purchased as crRNA from IDT. Two guides were designed to target exon 2 of the Khk gene, while the third targeted exon 4. The CRISPR/Cas9 system was delivered to embryos as a ribonucleoprotein (RNP) complex, which was formed using the protocol by Quadros et al. ^74^ with slight modifications. Briefly, 1ug of CRISPR-RNA (crRNA) containing the guide RNA sequence and 2µg of CRISPR transactivating RNA (tracrRNA) were annealed for each guide RNA, separately. Next, equi volume annealed crRNA and tracrRNA (ctRNA) representing the 3 guide sequences were assembled into an RNP complex with Alt-R® S.p. Cas9 Nuclease 3NLS recombinant protein. The complexes were cleared by centrifugation at 13,000 RPM 8 min and stored on ice until time of injection.

Microinjection and implantation: The ctRNA ribonucleoprotein (RNP) complexes were microinjected into the cytoplasm of CD-1:C57BL/6J hybrid embryos, resulting in 204 surviving embryos implanted into 5 pseudo-pregnant CD-1 females. This resulted in 13 live pups for genotyping.

Genotyping: Tail snips were collected 7-10 days after birth and genomic DNA extracted using Direct Tail PCR (Viagen) and Proteinase K (Qiagen). Genotyping was performed by PCR amplification of the genomic region surrounding the guide RNA recognition site and subsequent Sanger sequencing was performed. Primers are listed below. Four founder lines were generated, two of which were heritable (able to be passed to F1 generations). These mice display normal phenotypes in general when fed with a regular chow diet.

PCR primers:

Forward: GAAGTGTAGTCTCAGCAAGAG Reverse: GACACAACCTGAAGACTAAGTC

Founder lines: The third guide led to the deletion of a C in exon 4, which results in early termination codon (ETC) in exon 5. The founder was backcrossed into C57BL/6J background for at least five generations prior to phenotypic analyses.

#### Khk^+/+^/ODD-luc and Khk^-/-^/ODD-luc mice

*Khk^-/-^* mice and WT littermates (*Khk^+/+^*) were crossed with ODD-*luc* mice, respectively, to generate *Khk^-/-^/*ODD-*luc* and *Khk^+/+^*/ODD-*luc* mice. Successful expression of ODD-*luc* in *KHK*^-/-^ and *Khk^+/+^* mice were confirmed by ODD-*luc* transgene genotyping and whole-body bioluminescence imaging (BLI).

#### *In vivo and Ex vivo* bioluminescence imaging (BLI)

For *in vivo* BLI, ODD-*luc* mice were IP injected with D-luciferin potassium salt at the dose of 150mg/kg BW. Fifteen minutes later, mice were placed in the chamber and BLI was performed by using PhotonIMAGER^TM^ OPTIMA (Biospace Lab, Paris, France) under anesthesia with isoflurane aspiration (2%, 1000cc/circuit flow). For *ex vivo* BLI, ODD-*luc* mice were IP injected with D-luciferin potassium salt at the dose of 150mg/kg BW 5mins before euthanasia. The full-length GI tracts (from stomach to anus) were surgically removed after euthanasia by cervical dislocation. The first image was taken in approximately 15 min post D-luciferin injection and continued every 5 mins for 45mins. Photons were collected for a period of 60s, and images were obtained using Photo Acquisition software (Biospace lab, Paris, France), and analyzed using M3 Vision software (Biospace lab, Paris, France). Color bar indicates photons/(ph/s/cm2/sr) with minimum and maximum threshold values ^22^. Luciferase activity was calculated by the area under the kinetic curve (AUC) of luminescence.

#### Chronic sugar feeding on intestinal HIF-α stability in ODD-*luc* mice

Age-matched adult male and female ODD-*luc* mice were fed with rodent chow diet and free access to tap water or tap water containing 15% or 30% fructose (w/v), 15% or 30% glucose (w/v) or a 1:1 mixture of fructose:glucose for 2 weeks. *Ex vivo* BLI was performed to evaluate intestinal HIF-α luciferase activity at the end of the experiment.

#### Acute sugar gavage on intestinal HIF-α stability in ODD-*luc* mice

Age-matched adult male and female ODD-*luc* mice were fed with rodent chow diet. Mice received with daily gavage of saline or 20% fructose (w/v) or 20% glucose (w/v) solved in saline or a 1:1 mixture of fructose:glucose at the dose of 2 g/kg for 3 consecutive days. *Ex vivo* BLI was performed to evaluate intestinal HIF-α luciferase activity 15 minutes after the last dose of sugar gavage.

#### Streptozotocin (STZ)-induced hyperglycemia on intestinal HIF-α stability

Adult ODD-*luc* or C57BL/6J mice were IP injected with STZ at the dose of 100mg/kg BW for two consecutive days. Blood glucose measurements were performed using an ACCU-CHEK Aviva glucose meter and test strips. Hyperglycemic ODD-*luc* mice were used for intestinal *ex vivo* BLI two weeks after STZ injection. ^27^ C57BL/6J mice received pimonidazole (PMDZ) IP injection at the dose of 60mg/kg one hour before euthanasia.

#### Aldose reductase (AR) inhibition on intestinal HIF-α stability in ODD-*luc* mice

Eight-week-old male ODD-*luc* mice were fed with rodent chow diet and had free access to tap water. Mice received zopolrestat or saline via IP injection once daily at the dose of 50mg/kg BW ^31^ followed by daily gavage of glucose at a dose of 2g/kg BW for 3 days. *Ex vivo* BLI was performed on day 3, 15-20 minutes after final gavage of glucose to evaluate intestinal HIF-α stability.

#### Oral Sugar tolerance test

Ten-to twelve-week-old ODD-*luc* mice were fed normal chow followed by daily gavage of fructose (F), glucose (G) or fructose + glucose (FG) at a 1:1 mixture (all at the dose of 2 g/kg) for 3 days. The entire GI tract was then removed for BLI 15 minutes after last dose. Mice were orally gavaged with 20% sugars (glucose, fructose, or 1:1 mixture of glucose and fructose in saline) at the dose of 2 g/kg body weight after five hours fasting. Blood samples were obtained by tail nick and glucose levels were measured at 0, 15, 30, 60, and 120 min using an ACCU-CHEK Aviva glucose meter and test strips.

#### KHK inhibitor on intestinal HIF-α stability in ODD-*luc* mice

Adult male and female ODD-*luc* mice were fed with regular chow diet and 30% (w/v) fructose (F) via drinking water for 2 weeks *ad libitum*. KHK inhibitor (KI) or vehicle control (V) was administered twice daily via oral gavage at the dose of 10 mg/kg for the last 10 days. ^75^ *Ex vivo* BLI was performed to evaluate intestinal HIF-α luciferase activity one hour after the last dose of KI gavage.

#### PKM2 inhibitor on intestinal HIF-α stability in *Khk^-/-^*/ODD-*luc* mice

A selective PKM2 inhibitor ^39,40^, shikonin or vehicle control (DMSO) were IP injected to male adult *Khk^-/-^*/ODD-*luc* mice at the dose of 5mg/kg daily for three days. The entire GI tract BLI was performed 2 hours after the last dose of shikonin.

#### Fructose feeding on iron homeostasis

Four-month-old male *Khk^-/-^* mice and WT littermates were fed with chow diet and 30% fructose (w/v) *ad libitum* via drinking water for 2 weeks. Mice received PMDZ IP injection at the dose of 60mg/kg one hour before euthanasia. Plasma iron was measured at the end of the experiment. Complete blood count (CBC) was analyzed using VetScan HM5 hematology analyzer. Liver and small intestine were harvested for RNA-seq, RT-PCR and PMDZ staining.

#### Neutrophil mobilization

Age-matched 3-4-months-old male and female *Khk^-/-^* mice and WT littermates were injected subcutaneously with AMD3100 (CXCR4 antagonist) at the dose of 5mg/kg BW and CBC were analyzed by VetScan HM5 hematology analyzer 2 hours after injection. ^50^

#### Low iron diet

Four- to six-week-old male and female *Khk^-/-^* mice and WT littermates were fed with iron-replete AIN93G diet containing 350 ppm of iron or iron-deficient AIN93G diet (less than 5 ppm of iron) for 3 weeks. ^18^

#### Phz-induced hemolysis

Five-month-old male *Khk*^-/-^/ODD-*luc* mice were IP injected with phenylhydrazine (Phz) at the dose of 60mg/kg daily for 2 consecutive days. The entire GI *ex vivo* BLI were performed 48 hours after the last dose of Phz. ^18,37^

#### STAM HCC model

Pups were IP injected with 200 µg of STZ in 0.1 M sodium citrate buffer (pH 4.5) within 5 days after birth, followed by feeding with 60% high-fat diet (HF) *ad libitum* from the age of 3 weeks for 20 weeks. ^52,76^ Sugar water started [10% glucose (w/v), 30% fructose (w/v), or 30% fructose (w/v) + DFP] ^8^ after 6 weeks of HF (week 9-23, a total of 14 weeks). Deferiprone (DFP) was orally administered at the dose of 0.075mg/g BW via drinking water. ^61^

#### DEN/HF HCC model

Pups at 15-day-old were IP injected with N-Nitrosodiethylamine (DEN) at the dose of 25mg/kg in saline, followed by feeding with 60% high-fat diet (HF) *ad libitum* from the age of 6 weeks for 28 weeks. ^11,77^

#### Intestine-specific *Dmt1* knockout mice (*Dmt1* ^ΔIEC^)

*Dmt1* ^ΔIEC^ mice were generated by crossing *Dmt1*^fl/fl^ mice with *Dmt1*^fl/fl;Vil-cre-ERT^^2^ mice. Male *Dmt1*^fl/fl^ and *Dmt1* ^ΔIEC^ were subjected to the DEN/HF HCC model with additional 15% fructose (w/v) *ad libitum* via drinking water. Mice were IP injected with tamoxifen solution at the dose of 100mg/kg BW, daily for 3 consecutive days after 5 weeks of HF. Homozygous flox mice served as controls and received equal amount tamoxifen injection. After one week of tamoxifen washout, mice were fed with fructose drinking water for 22 weeks until the end of DEN/HF HCC model. To make a tamoxifen solution, forty microgram tamoxifen was suspended in 100 μl ethanol and solved in 900 μl corn oil (40mg/ml) by shaking rigorously at 55°C.

#### Bone marrow cell isolation, staining and flow cytometry analysis

Bone marrow cells were harvested from femurs and tibias of age-matched 5-6-month-old male *Khk^-/-^* mice and WT littermates as described.^78^ Cell suspensions were filtered through a 40-μm cell strainer. Cell number was counted using a hemocytometer method. For analysis of surface markers, cells were stained in RPMI 1640 medium containing 1% (wt/vol) BSA. The following antibodies were used at a dilution of 1/100–1/200: CD45-FITC, CD11b-PerCp-CY5.5, Ly6G-PE. Flow cytometry data were acquired on the FACS Carton II (Becton Dickinson) and analyzed using FlowJo 10.10.1 software (Treestar).

#### Isolation of mouse intestinal crypts and growth of intestinal organoids

Intestinal crypts were isolated and organoids were cultured using IntestiCult Organoid growth medium follow supplier’s instruction, as described. ^25,79^ Crypts were counted under the microscope.

1,500 crypts were resuspended in 300 µL of InstestiCult Organoid growth Medium, mixed (1:1) with Corning Matrigel matrix, and seeded onto 24-well plates with 50µl/well, covered with 750 µl IntestiCult Organoid Growth Medium. The growth medium was changed every two or three days. On day 6, the intestinal organoids were transferred to Seahorse XFe96/XF Pro cell culture microplates (Agilent Technologies, Inc., Santa Clara, CA) for Seahorse analysis.

#### Seahorse Analysis

20-30 organoids in 3 µl of Matrigel® per well were seeded onto XFe96 microplate and covered with 80 µl IntestiCult Organoid Growth Medium for 72 hours. Then, the organoids were treated with inhibitors, including Compound 968 (10µM), AOA (1mM), R162 (40µM), UK-5099 (10µM), and DCA (20mM) 16 hours before the assay. One hour before the measurements, culture medium was replaced, and the plate was incubated for 60 min at 37 °C in CO_2_ free incubator. For the mitochondrial stress test, culture medium was replaced by Seahorse XF DMEM medium (sugar free), supplemented with 150µM pyruvate, with or without 2mM glutamine. Then, standard mitochondrial stress test was performed on Seahorse XF96 analyzer (Agilent Technologies, Inc., Santa Clara, CA). ^80^ Oxygen consumption rates (OCR) were measured on injection of 15 mM glucose or fructose or 7.5mM glucose/7.5mM fructose (Port A), 5 µm oligomycin (Port B), 2 µM FCCP (Port C), and 2 µM Rotenone/2 µM Antimycin A (Port D) at 14, 34, 65 and 85 min, respectively. The first measurements after sugar injections were preceded by 4 min mixture time, followed by 0 min waiting time. The measurements after oligomycin injections were preceded by 5 min mixture time, followed by 10 min waiting time. OCR values were normalized to the total amount of protein content per well. ^80^

#### Liver enzymes and plasma biochemical assay

Liver enzymes and plasma biochemical assays were performed with commercially available kits: alanine aminotransferase (ALT), aspartate aminotrans-ferase (AST); Plasma iron was measured by Pointe iron reagent kit; Plasma erythropoietin (EPO) and hepcidin were measured by ELISA kit following the manufacturer’s instructions.

#### H&E staining

Formalin-fixed, paraffin-embedded liver sections were cut at 5µm thickness and stained with hematoxylin and eosin (H&E).

#### Perls’ Prussian blue iron staining

The distribution of storage iron in deparaffinized formalin-fixed tissues sections was visualized by using Perls’ Prussian blue staining and nuclear fast red counter stain. ^51,81^

#### PMDZ staining

PMDZ was IP injected at 60mg/kg BW one hour before euthanasia. Duodenum, Jejunum, ileum, and colon were fixed in 10% formalin and embedded with paraffin. Sections were stained with PMDZ (red) and counterstained with DAPI (blue) using Hypoxyprobe™ Kit follow manufacture’s instruction.

#### Bulk RNA-seq

Liver and intestine RNA were extracted using Qiagen RNeasy Mini kit following manufacturer’s instruction. RNA integrity was tested by Agilent 2100 bioanalyzer with RNA Integrity Number (RIN) between 7-9. Poly A enriched mRNA was used for library construction to avoid genomic DNA contamination. Constructed sequencing libraries were subject to sequencing using the Illumina Novaseq PE 150 platform (paired-end, 150 bp). Raw data (raw reads) of fastq format were first processed through in-house perl scripts. In this step, clean data (clean reads) were obtained by removing reads containing adapters, reads containing ploy-N and low-quality reads from raw data. Index of the reference genome was built using Hisat2 v2.0.5 and paired-end clean reads were aligned to the reference genome (ensembl_mus_musculus_grcm38_p6_gca_000001635_8) using Hisat2 v2.0.5. Feature Counts v1.5.0-p3 was used to count the reads numbers mapped to each gene and then fragments per kilobase of transcript per million mapped reads (FPKM) of each gene was calculated based on the length of the gene and reads count mapped to this gene. Differential expression analysis of two conditions/groups (two biological replicates per condition) was performed using the DESeq2 R package (1.20.0). Genes with an adjusted P-value ≤0.05 found by DESeq2 were assigned as differentially expressed. Gene Ontology (GO) and Kyoto Encyclopedia of Genes and Genomes (KEGG) enrichment analysis of differentially expressed genes were implemented by the clusterProfiler R package. GO terms with corrected P values less than 0.05 were considered significantly enriched by differential expressed genes.

#### Quantitative real time PCR (qRT-PCR)

Hepatic and intestinal RNA were extracted using Qiagen RNeasy Mini Kit and quantified using a NanoDrop One UV–Vis spectrophotometer (ThermoFisher). cDNA was prepared using 400 ng total RNA by reverse transcription–PCR (RT–PCR) using qScript cDNA SuperMix according to the manufacturer’s instructions. qRT-PCR was performed on a QuantStudio 3 real-time PCR systems using Power SYBR™ Green PCR Master Mix, cDNA and specific primers. Primers were designed by using Primer-BLAST. The relative gene expression was analyzed using the 2 ^-ΔΔCt^ method ^82^ and normalized to the house keeping genes. All fold changes are normalized to the control.

#### Western blot

Liver, intestine and kidney tissues were homogenized in TPER buffer containing 1×Protease Inhibitor Cocktail (ThermoFisher). Protein concentrations of the lysates were determined by using the BCA Protein Assay. Equal amounts of protein were loaded and resolved on 4%-15% SDS-polyacrylamide gels and transferred to PVDF membrane. The membrane was blocked for 1 hour using 5% nonfat dry milk in Tris-buffered saline-Tween 20 (TBS-T) and probed with primary antibody (1:1000 dilution) for KHK, β-actin, overnight at 4°C, and incubated with the corresponding horseradish peroxidase-conjugated secondary antibody. Protein signals were visualized on ChemiDoc Imaging System (BIO-RAD) by using enhanced chemiluminescence (SuperSignal West Pico, Pierce).

### QUANTIFICATION AND STATISTICAL ANALYSIS

Data are presented as the mean ± SEM. The sample size (n) for each experiment is indicated in the corresponding figure legends. For comparison of two groups, significance was determined using unpaired, two-tailed Student’s t test or Mann-Whitney test. For comparison of more than two groups a one- or two-way ANOVA was performed as appropriately followed by Tukey’s multiple comparisons test. All statistical analyses were performed using Prism 10.6.1(892). A p-value of less than 0.05 was taken to be statistically significant.

