## SUPPLEMENTAL INFORMATION for "Dietary fructose promotes MASH/HCC progression through enhanced intestinal HIF-2α-dependent iron absorption"

**Figure S1. Related to Figure 1; Fructose aberrantly stabilizes intestinal HIF-α**
(A) Schematic experimental design for chronic sugar feeding (15%, w/v) on intestinal HIF-α stability. (B) Schematic workflow of *ex vivo* bioluminescent imaging (BLI). (C) Representative images of *ex vivo* BLI in male 14-week-old ODD-*luc* mice with chronic sugar feeding (15%, w/v). (D) Intestinal kinetic curve of measured HIF-α luciferase activity in terms of luminescence ($ph/s/cm^2/sr \times 30min$). (E) Calculated area under curve (AUC) of (D) (n=4-5). (F) Blood glucose level. (G) Schematic experimental design for chronic sugar feeding (30%, w/v) on intestinal HIF-α stability. (H) Representative images of *ex vivo* BLI in 8-week-old male and female ODD-*luc* mice with chronic sugar feeding (30%, w/v). (I) Intestinal kinetic curve of measured HIF-α luciferase activity in terms of luminescence ($ph/s/cm^2/sr \times 30min$). (J) Calculated area under curve (AUC) of (I) (n=3-5). (K) Sugar oral gavage on intestinal HIF-α stability in female adult ODD-luc mice (n=3-4). (L, M) Sugar oral gavage on intestinal HIF-α stability of 8-month-old male and female ODD-luc mice (n=5-7). (N) Schematic experimental design for STZ-induced hyperglycemia model. (O) Representative images of pimonidazole (PMDZ) IHC staining of 12-14-weeks-old male C57BL/6J mice (n=2). Scale bar, 50μm. (P) Blood glucose level (n=4). (Q) Schematic experimental design for endogenous fructose on intestinal HIF-α stability. (R) Representative images of *ex vivo* BLI in 8-week-old male and female ODD-*luc* mice treated with zopolrestat in the presence of daily glucose gavage. (S) Intestinal kinetic curve of measured HIF-α luciferase activity in terms of luminescence ($ph/s/cm^2/sr \times 30min$). (T) Calculated area under curve (AUC) of (S) (n=5-6). Males are designated as triangles, and females are designated as circles (M, T). Data are presented as the mean ± SEM, *$p<0.05$, one-way ANOVA followed by Tukey's multiple comparison test or unpaired t test. F, fructose; G, glucose; FG, fructose + glucose at a 1:1 mixture; C, control (tap water or saline as indicated); Veh, vehicle; STZ, Streptozotocin; S, saline; Z, zopolrestat.

**Figure S2. Related to Figure 2; Generation of *Khk*$^{-/-}$ mice and *Khk*$^{-/-}$/ODD-luc mice**
(A) Cecum/BW ratio and blood glucose levels of nonfasted and fasted ODD-*luc* mice related to Figure. 2B (n=3-5). (B) DNA sequencing analysis of RT-PCR product of founder mice. The target exon 4 of khk was amplified using PCR. The top black line is "wild type". The short line underneath is the guide location. The chromatogram shows a heterozygous deletion (middle) and a homozygous deletion of a C in *Khk* gene (bottom). (C) Representative whole-body BLI images of male *Khk*$^{+/+}$/ODD-*luc* and *Khk*$^{-/-}$/ODD-*luc* mice. Data are presented as the mean ± SEM. * $p<0.05$, unpaired t test.

**Figure S3. Related to Figure 3; KHK deficiency prevents fructose-induced increases in plasma iron levels while disrupting systemic iron homeostasis**
(A) Representative images of duodenum PMDZ IHC staining related to Figure. 3A-C (n=2-4). Scale bar, 500μm. (B) Gating strategy of flow cytometry analysis of bone marrow cells. (C) Representative images and (D) Quantitative analysis of $CD45^+CD11B^+Ly6G^+$ bone marrow cells in 5-6-month-old male mice by flow cytometry (n=4). Data are presented as the mean ± SEM. * $p<0.05$, unpaired t test.

**Figure S4. Related to Figure 4; KHK deficiency inhibits iron absorption in a HIF-2α-dependent manner**
(A) RT-qPCR analysis of HIF-2α target genes in duodenal samples (n= 5-6). (B) Plasma hepcidin (n=6). (A) and (B) are related to Figure. 4A-C. Males are designated as triangles, and females are designated as circles. (C) Representative images of gross morphology (left, gross picture of euthanasia) (right, blood after centrifugation) of Phz-induced hemolytic anemia model. (D) Perls' Prussian blue iron staining of spleen section of *Khk*$^{+/+}$/ODD-*luc* and *Khk*$^{-/-}$/ODD-*luc* mice. (C) and (D) are related to Figure. 4D-F (n=4-5). Scale bar, 100μm. Data are presented as the mean ± SEM. * $p<0.05$, two-way ANOVA followed by Tukey's multiple comparison test. Phz, phenylhydrazine.

**Figure S5. Related to Figure 5; The KHK/HIF axis plays a critical role in MASH/HCC progression**
(A) Body weight growth trajectory, and (B) Calorie intake related to Figure. 5A-D. (C) Body weight growth trajectory, and (D) Calorie intake related to Figure. 5E-G. (E) Body weight growth trajectory, and (F) Calorie intake related to Figure. 5H-K. (G) Intestine HIF-α luciferase activity of male and female *Khk*$^{+/+}$/ODD-*luc* mice in response to STAM model for 8 weeks by BLI (n=8). (H) Liver weight and liver/body weight ratio (n=8). (I) Representative images of liver H&E staining. Scale bar, 100μm. Data are presented as the mean ± SEM. * $p<0.05$, unpaired t test.

**Figure S6. Related to Figure 6; Fructose promotes MASH/HCC progression in an iron-dependent**

**manner**

(A) Body weight growth trajectory, (B) Calorie intake, and (C) Complete blood count (CBC) analysis related to Figure. 6A-E (n=7-9). (D-K) Male *Dmt1$^{F/F}$ and Dmt1$^{\Delta IEC}$* mice subjected to DEN/HF HCC model. (D) Schematic timeline of intestine-specific *Dmt1* deletion on MASH/HCC progression. (E) Body weight growth trajectory. (F) Calorie intake. (G) Representative images of gross morphology of mice (left) and blood after centrifugation (right). (H) Blood HGB, HCT, RBC, MCV, MCH and MCHC (n=5-6). (I) Representative images of spleen morphology (top) and spleen weight, spleen/body weight (SW/BW) ratio and blood PLT (bottom) (n=5). Scale bar, 1cm. (J) Representative gross morphology of mice (top panel) and liver (bottom panel). Scale bar, 1cm. (K) HCC development (n=5). Data are presented as the mean ± SEM. * $p<0.05$, unpaired t test. G, glucose; F, fructose; DFP, deferiprone. WBC, total white blood cell count; PLT, platelets; HF, high-fat diet; TAM, tamoxifen; HGB, hemoglobin; RBC, red blood cell; HCT, hematocrit; MCH, mean corpuscular hemoglobin; MCV, mean corpuscular volume; MCHC, mean corpuscular hemoglobin concentration. All cartoons were created in Biorender.com.

**Table S1. Related to Figure 2-4**; Primers used for quantitative RT-PCR.

| | |
|---|---|
| *Hif1a* Forward | ATAGCTTCGCAGAATGCTCAGA |
| *Hif1a* Reverse | CAGTCACCTGGTTGCTGCAA |
| *Ldha* Forward | TGTCTCCAGCAAAGACTACTGT |
| *Ldha* Reverse | GACTGTACTTGACAATGTTGGGA |
| *Eno1* Forward | TGCGTCCACTGGCATCTAC |
| *Eno1* Reverse | CAGAGCAGGCGCAATAGTTTTA |
| *Pdk1* Forward | TTACTCAGTGGAACACCGCC |
| *Pdk1* Reverse | GTTTATCCCCCGATTCAGGT |
| *Bnip3* Forward | GTTACCCACGAACCCCACTTT |
| *Bnip3* Reverse | GTGGACAGCAAGGCGAGAAT |
| *Vegfa* Forward | CTGCTGTAACGATGAAGCCCTG |
| *Vegfa* Reverse | GCTGTAGGAAGCTCATCTCTCC |
| *Hif2a exon2* Forward[1] | TGAGTTGGCTCATGAGTTGC |
| *Hif2a exon2* Reverse[1] | TATGTGTCCGAAGGAAGCTG |
| *Epo* Forward | GGTACTGGGAGCTCAGGAATTG |
| *Epo* Reverse | TGTGAGTGTTCGGAGTGGAG |
| *Dmt1 +IRE* Forward[1] | TGTTTGATTGCATTGGGTCTG |
| *Dmt1 +IRE* Reverse[1] | CGCTCAGCAGGACTTTCGAG |
| *Cybrd1* Forward | CATCCTCGCCATCATCTC |
| *Cybrd1* Reverse | GGCATTGCCTCCATTTAGCTG |
| *Slc40a1* Forward | ATGGGAACTGTGGCCTTCAC |
| *Slc40a1* Reverse | TCCAGGCATGAATACGGAGA |
| *Neu3* Forward | ATGGAGGCCACATTACCTGG |
| *Neu3* Reverse | TCTGGCACCTCTCAGTAACAT |
| *Hamp* Forward | TCTTCTGCATTGGTATCGCA |
| *Hamp* Reverse | GAGCAGCACCACCTATCTCC |
| *Fatp2* Forward | ACACACCGCAGAAACCAAATGACC |
| *Fatp2* Reverse | TGCCTTCAGTGGATGCGTAGAACT |
| *Fatp4* Forward | AGTAAGCATGTGGCTTTGGGCAAG |
| *Fatp4* Reverse | TTTGGCAGAAGATGGAGCAACAGC |
| *Cd36* Forward | AGATGACGTGGCAAAGAACAG |
| *Cd36* Reverse | CCTTGGCTAGATAACGAACTCTG |
| *Actb* Forward | TATTGGCAACGAGCGGTTCC |
| *Actb* Reverse | GGCATAGAGGTCTTTACGGATGT |

#### Figure S1

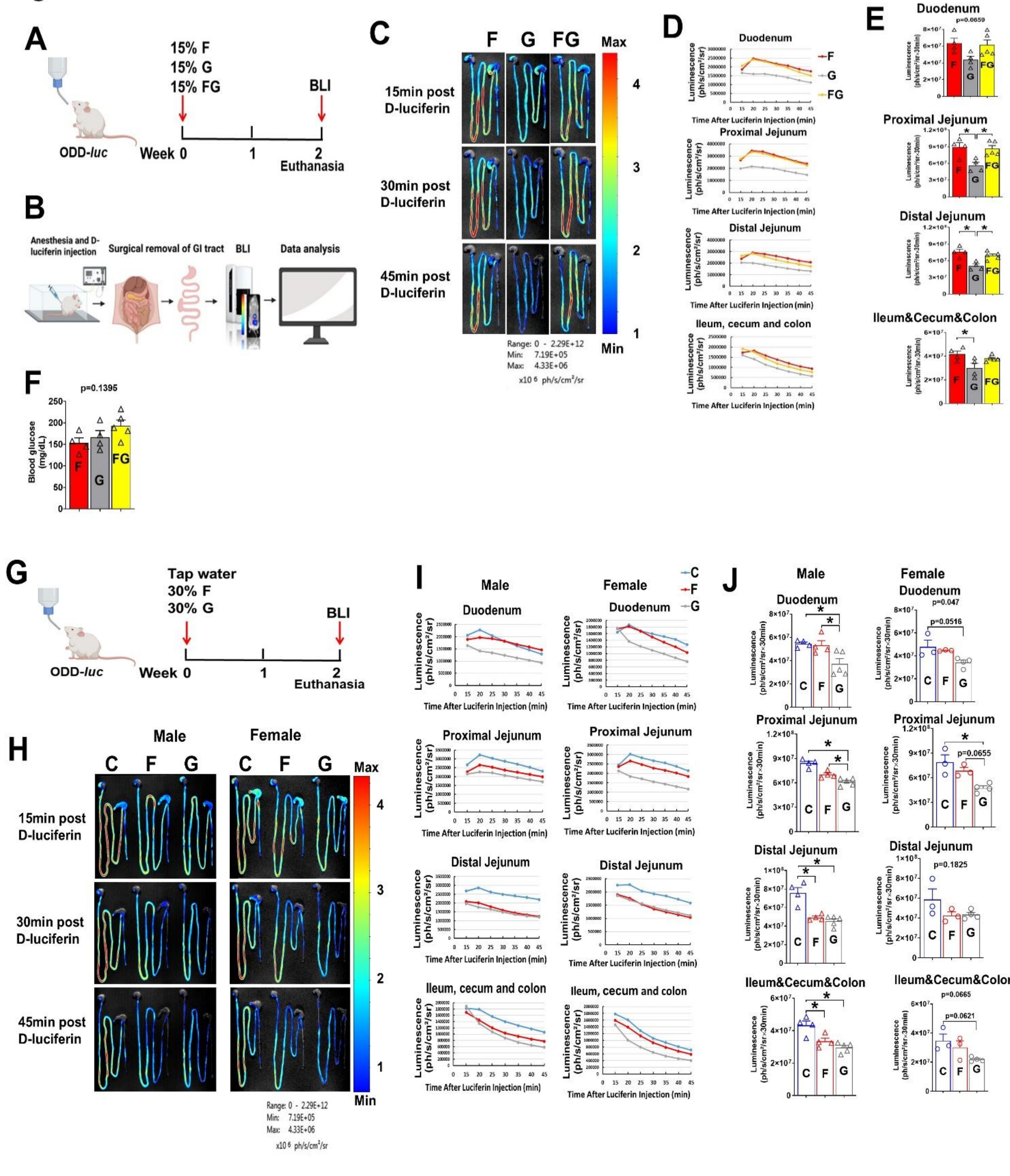

A
15% F
15% G
15% FG
BLI
ODD-luc
Week 0
1
2
Euthanasia
B
Anesthesia and D-luciferin injection
Surgical removal of GI tract
BLI
Data analysis
C
F
G
FG
Max
15min post D-luciferin
30min post D-luciferin
45min post D-luciferin
Min
Range: 0 - 2.29E+12
Min: 7.19E+05
Max: 4.33E+06
x10 6 ph/s/cm²/sr
D
Duodenum
Proximal Jejunum
Distal Jejunum
Ileum, cecum and colon
Luminescence (ph/s/cm²/sr)
Time After Luciferin Injection (min)
E
Duodenum
p=0.0659
Proximal Jejunum
Distal Jejunum
Ileum&Cecum&Colon
F
p=0.1395
Blood glucose (mg/dL)
G
Tap water
30% F
30% G
BLI
ODD-luc
Week 0
1
2
Euthanasia
H
Male
Female
C
F
G
I
Male
Female
J
Duodenum
p=0.047
p=0.0516
Proximal Jejunum
p=0.0655
Distal Jejunum
p=0.1825
Ileum&Cecum&Colon
p=0.0665
p=0.0621

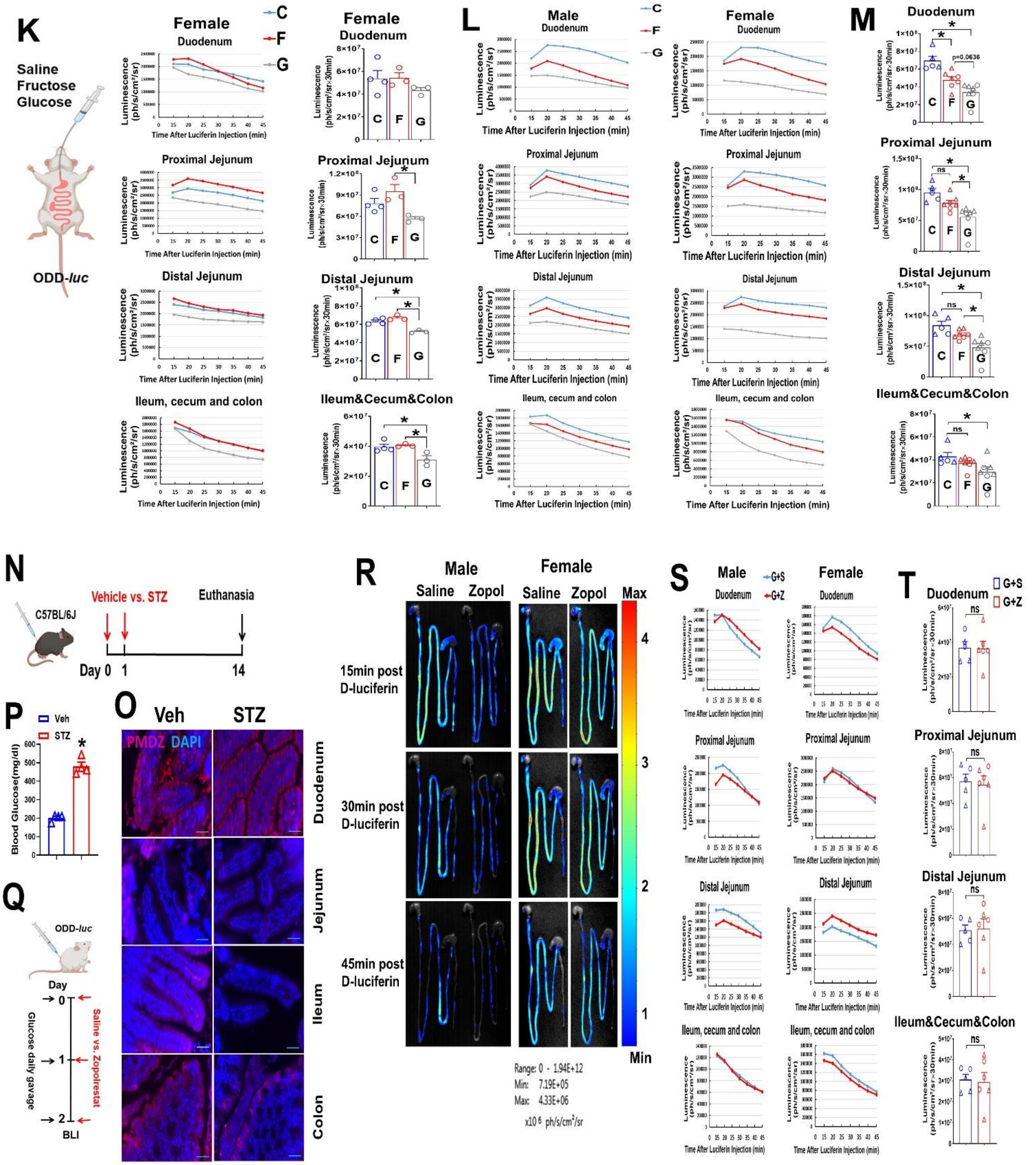
K
Saline
Fructose
Glucose
ODD-luc
Female
Duodenum
Proximal Jejunum
Distal Jejunum
Ileum, cecum and colon
Luminescence (ph/s/cm²/sr)
Time After Luciferin Injection (min)
C
F
G
Ileum&Cecum&Colon
L
Male
M
N
C57BL/6J
Vehicle vs. STZ
Euthanasia
Day 0 1 14
P
Veh
STZ
Blood Glucose(mg/dl)
O
PMDZ DAPI
Jejunum
Ileum
Colon
Q
Day
Glucose daily gavage
Saline vs. Zopolrestat
BLI
R
Saline
Zopol
15min post D-luciferin
30min post D-luciferin
45min post D-luciferin
Max
Min
Range: 0 - 1.94E+12
Min: 7.19E+05
Max: 4.33E+06
x10 6 ph/s/cm²/sr
S
G+S
G+Z
T
ns
p=0.0636

### Figure S2

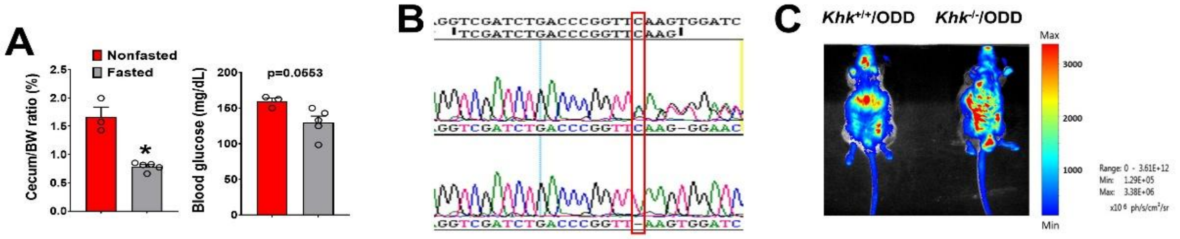


### Figure S3

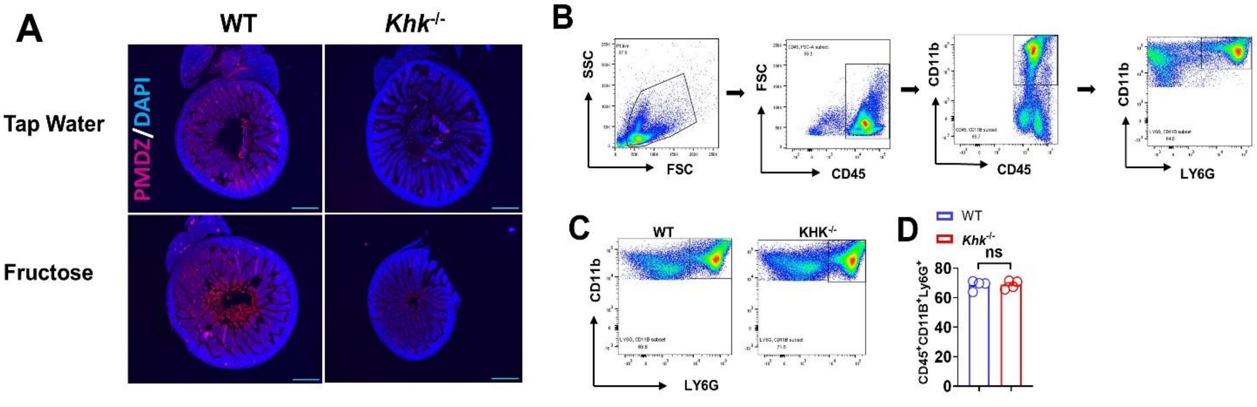


### Figure S4

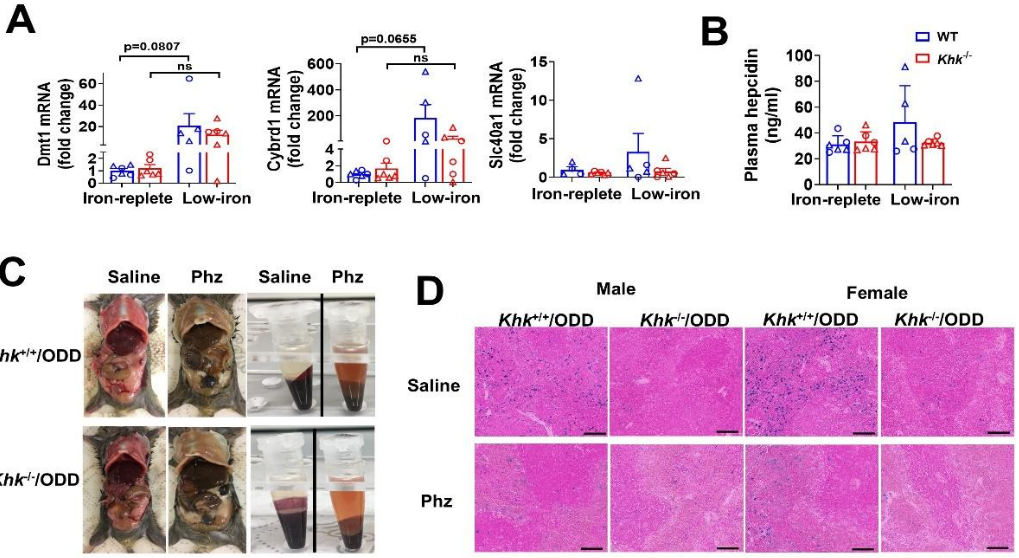

### Figure S5

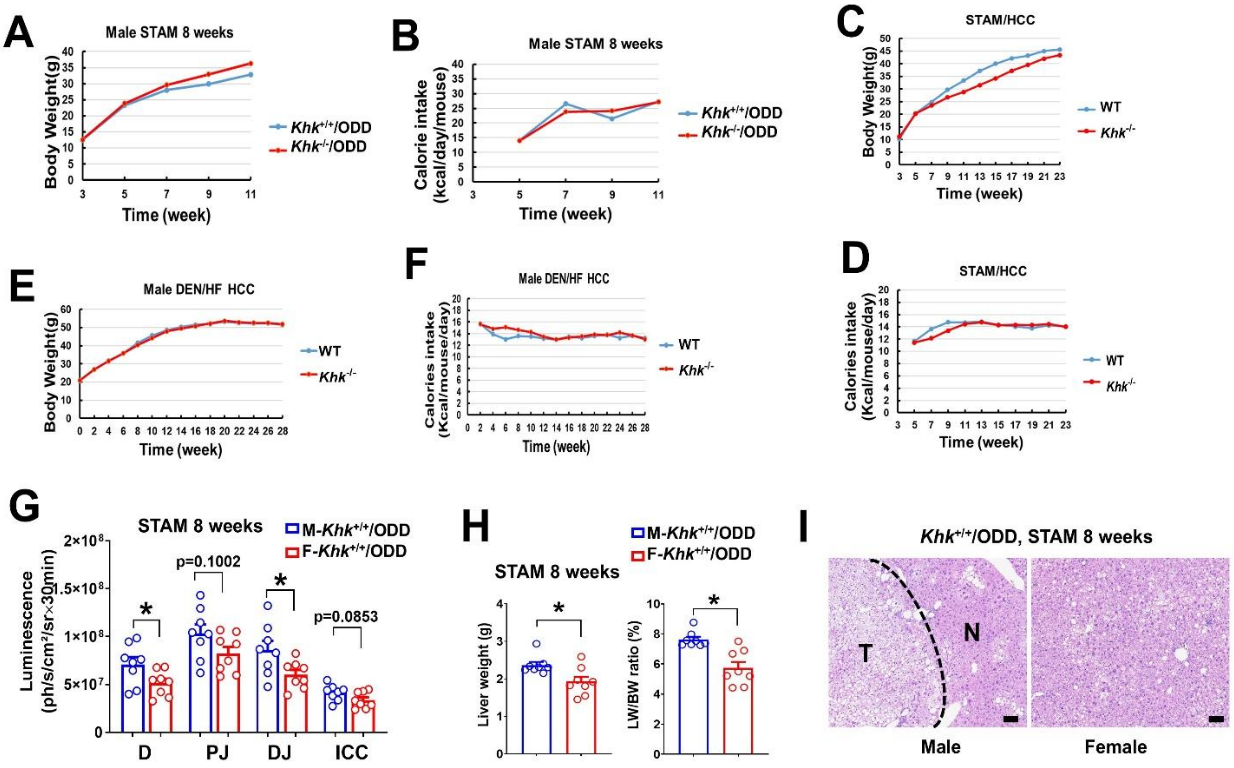
A
Male STAM 8 weeks
Body Weight(g)
Time (week)
Khk+/+/ODD
Khk-/-/ODD
B
Male STAM 8 weeks
Calorie intake (kcal/day/mouse)
Time (week)
Khk+/+/ODD
Khk-/-/ODD
C
STAM/HCC
Body Weight(g)
Time (week)
WT
Khk-/-
E
Male DEN/HF HCC
Body Weight(g)
Time (week)
WT
Khk-/-
F
Male DEN/HF HCC
Calories intake (Kcal/mouse/day)
Time (week)
WT
Khk-/-
D
STAM/HCC
Calories intake (Kcal/mouse/day)
Time (week)
WT
Khk-/-
G
STAM 8 weeks
M-Khk+/+/ODD
F-Khk+/+/ODD
Luminescence (ph/s/cm²/sr×30min)
p=0.1002
p=0.0853
D
PJ
DJ
ICC
H
STAM 8 weeks
M-Khk+/+/ODD
F-Khk+/+/ODD
Liver weight (g)
LW/BW ratio (%)
I
Khk+/+/ODD, STAM 8 weeks
T
N
Male
Female

### Figure S6

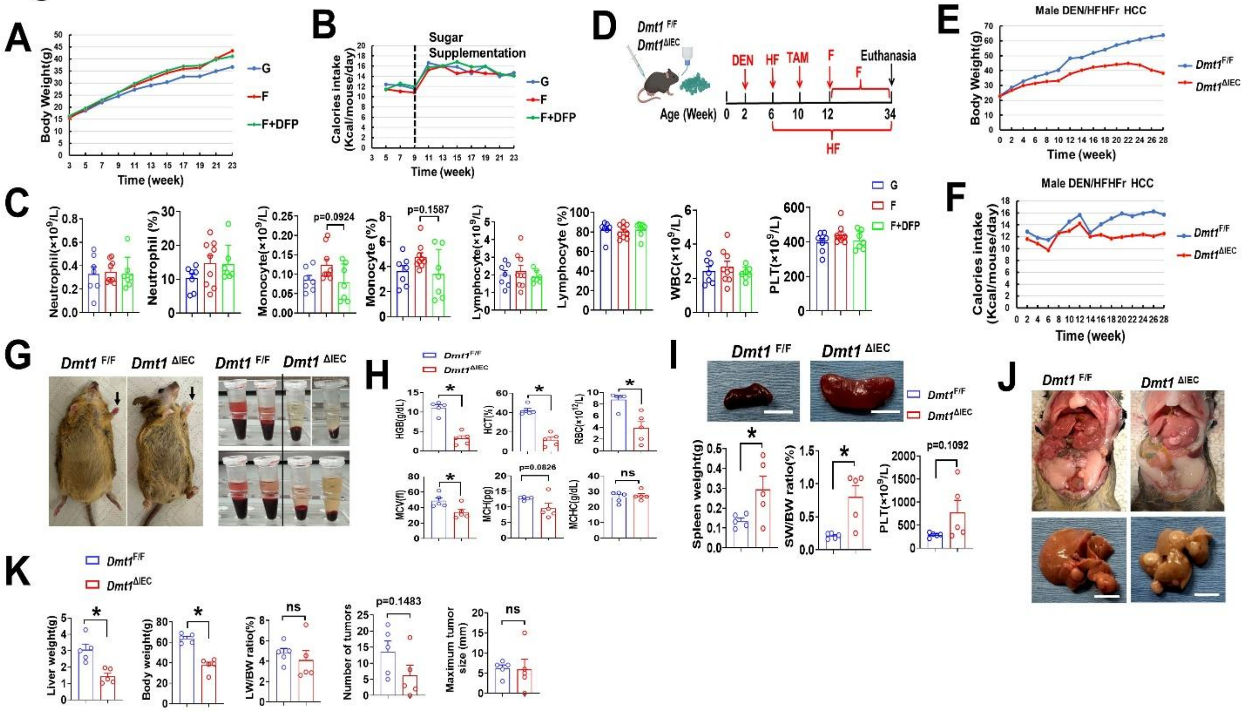

A
Body Weight(g)
Time (week)
G
F
F+DFP
B
Sugar Supplementation
Calories intake (Kcal/mouse/day)
Time (week)
D
Dmt1 F/F
Dmt1 ΔIEC
DEN
HF
TAM
F
Euthanasia
Age (Week)
HF
E
Male DEN/HFHFr HCC
Body Weight(g)
Time (week)
F
Male DEN/HFHFr HCC
Calories intake (Kcal/mouse/day)
Time (week)
C
Neutrophil(×10^9/L)
Neutrophil (%)
Monocyte(×10^9/L)
p=0.0924
Monocyte (%)
p=0.1587
Lymphocyte(×10^9/L)
Lymphocyte (%)
WBC(×10^9/L)
PLT(×10^9/L)
G
Dmt1 F/F
Dmt1 ΔIEC
H
HGB(g/dL)
HCT(%)
RBC(×10^12/L)
MCV(fl)
MCH(pg)
p=0.0826
MCHC(g/dL)
ns
I
Spleen weight(g)
SW/BW ratio(%)
PLT(×10^9/L)
p=0.1092
J
K
Liver weight(g)
Body weight(g)
LW/BW ratio(%)
ns
Number of tumors
p=0.1483
Maximum tumor size (mm)
ns
